# A minimal model of stable autocatalytic RNA-replication through integration of replication, molecular regulation, and energy conversion

**DOI:** 10.64898/2026.07.25.740712

**Authors:** Bernard Conrad, Christian Iseli, Magnus Pirovino

## Abstract

Catalysis and allostery are complementary principles of biological function: catalysis accelerates biochemical reactions, whereas allosteric regulation dynamically controls molecular interactions. The emergence and persistence of early autocatalytic ribozyme systems likely depended not only on template-directed self-replication but also on ATP production, utilization, and recycling through prebiotically plausible energy-conversion processes. We previously proposed that resistance to molecular parasitism in autocatalytic RNA networks can emerge through hyperparasitic regulation, whereby hyperparasitic ribozymes compete with parasitic ribozymes for binding to the host replicase while additionally binding parasitic ribozymes, thereby redirecting competitive interactions away from host exploitation. Here, we develop a minimal mathematical model of an autocatalytic RNA-world system in which replication, ATP/ADP-based energy conversion, and ATP-dependent allosteric regulation become dynamically coupled. Building on a general reaction framework describing ribozyme replication, catalytic interactions, and ATP/ADP cycling, we identify the minimal regulatory architecture required for stable RNA replication in the presence of parasitic mutants. Our simulations reveal a sequential evolutionary transition in which control precedes optimization. ATP-dependent allosteric regulation first evolves to tame molecular parasitism through hyperparasitic binding, whereby ATP-loaded parasitic ribozymes compete with ATP-free parasites for binding to the host replicase while also binding directly to parasitic ribozymes. This stabilizes RNA replication but simultaneously imposes an energetic cost by sequestering ATP and thereby reducing ATP–ADP turnover. The resulting regulatory burden creates selective pressure for the evolution of ATP synthase/ATPase ribozymes that accelerate ATP–ADP cycling and restore the metabolic flux required for sustained RNA replication. We further identify two plausible evolutionary routes to parasite control: parasitic ribozymes either become intrinsically allostery-prone or are converted into allostery-prone forms by an evolved allosterase ribozyme. Once ATP turnover is sufficiently rapid, both mechanisms confer long-term resistance to recurrent parasitic invasion. These results suggest that stable RNA-based evolution required the progressive integration of information replication, molecular reulation, and increasingly efficient metabolic energy conversion. More generally, the model identifies ATP-dependent allosteric regulation as a plausible evolutionary bridge linking autocatalytic RNA replication to the emergence of regulated proto-metabolism, transforming a parasite-limited replicating system into a self-regulating proto-biological organization capable of sustained evolutionary dynamics.

**Highlights:**

- A minimal autocatalytic RNA network achieves stable self-replication despite recurrent parasitic invasion
- ATP-dependent allostery tames molecular parasitism through hyperparasitic binding
- Control of molecular parasitism precedes optimization of metabolic energy conversion
- ATP sequestration creates an energetic burden that drives increased ATP–ADP turnover
- Stable parasite control evolves through either intrinsic allostery or allosterase-mediated regulation
- Stable RNA autocatalysis emerges through the sequential integration of replication, molecular regulation, and energy conversion

## 1. Introduction

Origin-of-life scenarios are traditionally framed around two contrasting principles: the metabolism-first and replication-first hypotheses (Anet, 2004; Pross, 2004). The metabolism-first view proposes that complex, self-organized reaction networks initially redirected energy from planetary processes into organic chemistry, with prebiotic geochemical catalysis gradually giving rise to biological catalysis (Zimmermann et al., 2024). In this framework, the architecture of ancient chemolithoautotrophic metabolisms may provide models of primordial nonenzymatic reaction networks, exemplified by carbon fixation pathways such as the reductive citric acid cycle (Morowitz, 1999).

Adenosine triphosphate (ATP) occupies a central position at the interface of metabolism and heredity. It serves as the universal energy currency of extant life and of the most ancient reconstructed autotrophic metabolisms based on H_2_ and CO_2_, while simultaneously constituting one of the four nucleotide building blocks of RNA (De Duve, 2005). Several core reactions of autotrophic metabolism can occur spontaneously in the absence of enzymes, suggesting that ATP may have arisen early as a product of protometabolism driven by H_2_ and CO_2_ (Pinna et al., 2022). However, purine biosynthesis requires multiple phosphorylation steps coupled to ATP hydrolysis, implying the existence of a prebiotic phosphorylating equivalent. A conserved metabolic intermediate, acetyl phosphate, can phosphorylate ADP to ATP in aqueous solution with yields approaching 20% in the presence of Fe^3+^ ions. Its relative specificity toward adenosine suggests that ATP synthesis may have been favored under prebiotic conditions, potentially explaining its universal conservation across the tree of life (Pinna et al., 2022).

Coenzymes — ATP itself formally belongs to this class — are small organic cofactors that bind specifically to enzymes and participate directly in biochemical transformations, being required by up to 30% of modern enzymes (Zimmermann et al., 2024). Several features suggest that coenzymes represent molecular relics of early biochemical systems. Many share structural similarities with RNA building blocks, particularly the adenosine monophosphate moiety; they appear to have undergone limited structural modification since the last universal common ancestor; and they retain their primary catalytic role in facilitating biochemical reactions (Kirschning, 2022).

These observations point to a close connection between early metabolic chemistry and RNA, widely considered the primordial carrier of genetic information and catalysis (Papastavrou, 2024; Gianni, 2026). According to Harold J. Morowitz (Zimmermann et al., 2024; Morowitz, 1999), the diversity of biomolecules radiating from a small core of carbon-fixation reactions suggests that the molecular precursors of genetic systems emerged as products of expanding protometabolic networks. This view contrasts with the alternative hypothesis that RNA itself initiated protometabolism through its catalytic capabilities, the central premise of the replication-first model.

Regardless of the sequence of events, catalysis is central to all origin-of-life scenarios, with catalytic activity modulated by cofactors such as metals, metal ions, and coenzymes (Zimmermann et al., 2024). Early biochemical systems must have rapidly acquired the capacity to adapt to environmental fluctuations. In this context, the discovery of allostery — defined as a change in ligand binding in the presence of a second ligand (Fenton, 2008) — was famously described by Jacques Monod as the “second secret of life,” second only to the genetic code (Monod, 1997). Modern mechanistic interpretations of allostery often adopt an ensemble perspective, derived from the Monod-Wyman-Changeux model (Wodak et al., 2019), in which allosteric behavior arises from the intrinsic free-energy landscape of a macromolecule and from perturbations such as ligand binding, protonation, or intermolecular interactions that reshape this landscape.

The RNA world hypothesis proposes that early evolving systems relied on RNA molecules capable of both storing genetic information and catalyzing chemical reactions (Gilbert, 1986). Ribozymes may have arisen spontaneously through abiotic polymerization of RNA oligomers. Random RNA sequences naturally generate abundant secondary structures, including hairpins, among which a fraction can display catalytic activities such as RNA ligation. Such activities could enable modular RNA evolution through recombination and fragment assembly (Briones et al., 2009). Indeed, Group I intron ribozymes possessing recombinase activity have been shown to self-assemble from inactive oligonucleotide fragments, acquiring the capacity for self-reproduction through autocatalysis (Hayden and Lehman, 2006; Kauffman, 1986). Progressive evolutionary refinement could subsequently have produced RNA polymerase ribozymes capable of template-directed replication, initiating a Darwinian process comparable to modern biological evolution (Nghe, 2025).

In previous work, we developed mathematical models of early RNA-based life autocatalytic systems inspired by the RNA-world hypothesis, focusing on the dynamics of molecular replication and parasitism (Conrad et al., 2023; Pirovino et al., 2025). Using a Lotka–Volterra framework, we first demonstrated that tripartite habitats composed of nested host–molecular parasite pairs can attain homeostatic stability (Conrad et al., 2023). We subsequently extended this approach to autocatalytic ribozyme networks governed by Michaelis–Menten kinetics, showing that evolutionary interactions among host, parasite, and hyperparasite catalysts generate characteristic trajectories in which parasite integration emerges as a distinctive dynamical feature of the network (Pirovino et al., 2025).

Here, we further extend this framework by coupling autocatalytic RNA replication to an ATP/ADP-based proto-metabolic energy cycle and ATP-dependent allosteric regulation. Within this generalized reaction network, we identify a minimal autocatalytic RNA system capable of sustained self-replication despite the continual emergence of non-catalytic parasitic mutants. The minimal architecture comprises a self-replicating host ribozyme *R*_1_, a parasitic ribozyme *R*_3_ that evolves catalytic activity for the ATP/ADP cycle, and ATP-dependent allosteric regulation of parasitic ribozymes mediated by a hyperparasitic binding mechanism, i.e. high affinity for the host replicase comparable to that of the parasite, high affinity for ATP-free parasitic ribozymes, and optionally high affinity for ATP-loaded parasitic ribozymes. Our simulations reveal a sequential evolutionary transition. An initial increase in ATP/ADP turnover, catalyzed by the metabolic ribozyme, establishes the energetic conditions required for ATP-dependent allosteric regulation to emerge. Once established, allosteric regulation suppresses parasite-driven collapse but imposes an energetic cost through ATP sequestration, necessitating a further increase in ATP/ADP turnover to sustain stable replication. Finally, we demonstrate two plausible evolutionary routes to stable parasite control: either parasitic ribozymes are intrinsically allostery-prone, or an allosterase ribozyme evolves that converts non-allostery-prone parasites into allostery-prone forms. Together, these results identify a minimal regulatory architecture in which progressively enhanced energy conversion, information replication, and molecular regulation become dynamically integrated, enabling robust autocatalytic evolution.

## 2. Methods

We consider an RNA-world habitat in which an underlying proto-metabolism provides sufficient chemical energy to sustain the production and maintenance of a limited set of ribozymes. These ribozymes are assumed to self-organize into autocatalytic networks.

### 2.1. Definitions for the ATP-regulated autocatalytic cycle model: the general model

In this section, we formulate the reaction equations of a general mathematical framework describing autocatalytic ribozyme networks fueled and regulated by an ATP/ADP-based energy cycle supplied by the surrounding habitat. The framework permits an arbitrary number of distinct ribozyme species participating in an autocatalytic replication network and provides the basis for the minimal model developed in the Results section.

#### 2.1.1 Ribozymes and Michaelis-Menten equations of three different types

Let *n* be the number of different ribozymes, *R*_1_, *R*_2_,…, *R*_*n*_, the free ribozyme concentrations, and 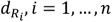, the decay rates of these ribozymes.

These ribozymes may exhibit three fundamentally distinct catalytic functions, which we refer to here as the three types of catalysis.

##### Type 1 catalysis – RNA modification

A catalytically active ribozyme modifies a substrate ribozyme to produce a different ribozyme. During this process, the catalyst forms a transient intermediate complex with the substrate ribozyme, which is chemically transformed into the reaction product.

##### Type 2 catalysis – template-directed replication

A catalytically active ribozyme uses another ribozyme as a template for the synthesis of a new ribozyme. The catalyst therefore forms a transient intermediate complex with the template ribozyme. Upon completion of replication, both the catalyst and the template are released unchanged, while the newly synthesized ribozyme enters the habitat as an additional product.

##### Type 3 catalysis – metabolic energy conversion

A catalytically active ribozyme catalyzes the interconversion of non-genetic substrates, such as ATP, ADP, or other small molecules involved in proto-metabolism. In contrast to Types 1 and 2, neither the substrates nor the products are information-carrying RNA molecules. Instead, these reactions generate or recycle the chemical energy and molecular resources required to sustain autocatalytic ribozyme replication.

In the present work, modeling of type-1 catalytic processes is restricted to hypothetical allosterases involved in allosteric ATP regulation of ribozymes. Although type-1 processes, including ligase- and recombinase-mediated RNA modification, were likely important during the emergence of template-based replication (type-2 catalysis), the present model focuses on the minimal autocatalytic pathway required to sustain replication. Within this framework, stable self-replication is achieved through the coupled action of type-2 and type-3 catalytic processes, while type-1 catalysis is considered only as a regulatory mechanism for controlling parasitic ribozymes.

##### Definitions of type-1 and type-2 catalytic processes

We first define the concentrations and rate processes associated with type-1 and type-2 ribozyme activities. Throughout the model, we use the general reaction scheme

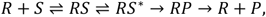

where *RS* denotes the reversibly bound ribozyme–substrate complex, *RS*^∗^ the activated catalytic complex, and *RP* the ribozyme–product complex. The activated catalytic complex *RS*^∗^ is explicitly represented in the model, whereas the ribozyme–product complex *RP* is not. Throughout this work, the term “charged” refers to the activated ribozyme–substrate state *RS*^∗^, whereas “uncharged” refers to the reversibly bound ribozyme–substrate complex *RS*.

Let *Y*′, *Z*′, *Y*′′, *Z*′′*ϵ* ℝ^nxn^ be matrices. For *i, j* = 1,.., *n* the entries of these matrices are defined as follows:

*Y*′_*ij*_ … uncharged type 1 intermediate complex of ribozyme *R*_*i*_ with substrate *R*_*j*_, being subject to modification,

*Z*′_*ij*_ … charged type 1 intermediate complex of ribozyme *R*_*i*_ with substrate *R*_*j*_, being subject to modification,

*Y*′′_*ij*_ … uncharged type 2 intermediate complex of ribozyme *R*_*i*_ with template *R*_*j*_,

*Z*′′_*ij*_ … uncharged type 2 intermediate complex of ribozyme *R*_*i*_ with template *R*_*j*_

We note that in general *Y*′_*ij*_ ≠ *Y*′_*ji*_, *Z*′_*ij*_ ≠ *Z*′_*ji*_, *Y*′′_*ij*_ ≠ *Y*′′ _*ji*_, and *Z*′′_*ij*_ ≠ *Z*′′_*ji*_. This stems from the fact that for both type 1 and type 2 intermediate complexes the roles of *R*_*i*_ and *R*_*j*_ are distinct, and thus not exchangeable: *R*_*i*_ acts as the enzyme catalyzing the reproduction, and *R*_*j*_ functions as type 1 substrate subject to modification or as type 2 template.

In both cases, type 1 and 2, these concentrations are governed by a respective Michaelis-Menten process:

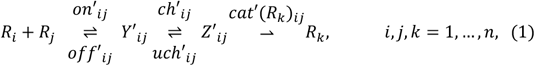

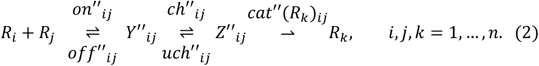

Here, the different classes of kinetic parameters—namely association (on) rates, dissociation (off) rates, charging (activation) rates, discharging/uncharging (deactivation) rates, and catalytic rates—are encoded as entries of *n* × *n* matrices. For instance, for *n* = 2, the type-1 association-rate matrix is defined as:

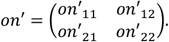

We write the *n* x *n* matrix *on*′ = [*on*′_*ij*_]_*ij*_ *ϵ* ℝ^nxn^, implying *i, j* = 1, …, *n*.

Thus, we define the following type-1 and-2 matrices of the rates:

*off*′ = [*off*′ _*ij*_]_*ij*_, *off*′′ = [*off*′′ _*ij*_]_*ij*_ *ϵ* ℝ^nxn^… off-rates from uncharged type 1 and 2 intermediate complexes to free ribozymes,

*ch*′ = [*ch*′ _*ij*_]_*ij*_, *ch*′′ = [*ch*′′ _*ij*_]_*ij*_ *ϵ* ℝ^nxn^… charge-rates from uncharged to the charged type 1 and 2 intermediate complexes,

*uch*′ = [*uch*′ _*ij*_]_*ij*_, *uch*′′ = [*uch*′′ _*ij*_]_*ij*_ *ϵ* ℝ^nxn^… uncharge-rates from the charged to uncharged type 1 and 2 intermediate complexes.

We define the catalytic rates as follows. Let *cat*′(*R*_1_) = [*cat*′(*R*_1_)_ij_]_*ij*_ ∈ ℝ^n×n^ denote the matrix of rates at which charged type-1 intermediate complexes catalyze the formation of ribozyme *R*_1_. More generally, we define

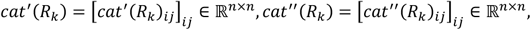

as the matrices of rates at which charged type-1 and type-2 intermediate complexes, respectively, catalyze the formation of ribozyme *R*_*k*_, *k* = 1, …, *n*.

Within this framework, each charged intermediate complex may catalyze the production of any free ribozyme species, either (i) through modification of an existing ribozyme in the case of type-1 catalysis, or (ii) through mutation or imperfect template-based replication in the case of type-2 catalysis.

Consequently, each free ribozyme species may receive contributions to its production from every charged intermediate complex. In practice, however, most entries *of cat*′(*R*_*k*_) and *cat*′′(*R*_*k*_) are expected to be zero or negligibly small for all *k, i, j* = 1, …, *n*.

Finally, let 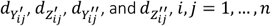, denote the decay rates of the corresponding catalytic intermediate complexes.

##### The ATP-ADP cycle and its type 3 catalytic process

We now introduce two further concentrations, *T* and *D*, responsible for the energy supply in the habitat providing the metabolism needed for ribozyme reproduction.

*T* … concentration of free ATP in the habitat, and

*D* … concentration of free ADP in the habitat.

These concentrations are governed by the following reaction equation:

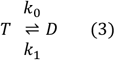

We assume that the total amount of ADP and ATP in the habitat remains constant, including both free and bound forms. This constraint defines the external metabolic conditions for the reproduction cycle of the ribozymes *R*_*i*_. The ATP–ADP cycle (3) is further assumed to be linked to the Michaelis–Menten processes (1) and (2) through two coupling pathways.

###### The first coupling between cycles (3), (1), and (2)

We assume that the catalytic Michaelis–Menten processes (1) and (2) are driven by the ATP–ADP cycle (3) as follows. The activation of intermediate complexes formed between two ribozymes *R*_*i*_ and *R*_*j*_, from the uncharged to the charged state, requires ATP consumption. For type 1 and type 2 processes, this activation is represented by:

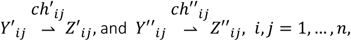

During these transitions, ATP is hydrolyzed to ADP. Consequently, the production of an additional amount 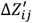 or 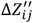 requires the consumption of a corresponding amount of ATP, denoted by Δ*T*′and Δ*T*′′, respectively. These quantities are determined by the following equations:

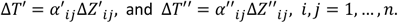

Here, the matrices 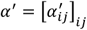 and 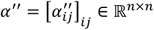 contain the coefficients specifying the ATP requirements of the charging processes described by reactions (1) and (2), respectively.

###### The second coupling between cycle (3) and processes (1) and (2)

We assume that some, or potentially all, ribozymes *R*_*i*_ can contribute to the catalysis of the ATP–ADP cycle by enhancing both the synthesis of ATP and its hydrolysis into ADP. Following the definition introduced above, these reactions are classified as type 3 catalytic processes:

*Y*′′′(*D*)_*i*_ … concentration of uncharged type 3 ribozyme-ADP intermediate complex,

*Z*′′′(*D*)_*i*_ … concentration of charged type 3 ribozyme-ADP intermediate complex,

*Y*′′′(*T*)_*i*_ … concentration of uncharged type 3 ribozyme-ATP intermediate complex,

*Z*′′′(*T*)_*i*_ … concentration of charged type 3 ribozyme-ATP intermediate complex, *i* = 1, …, *n*.

The catalysis of this metabolic cycle is governed by Michaelis–Menten kinetics in both the forward and reverse directions. Considering first the direction of ATP production:

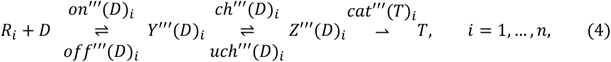

and second the direction of ATP-consumption and-hydrolysis, respectively:

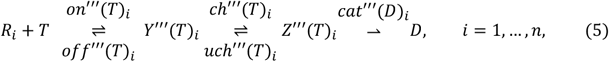

Here, the vectors *on*′′′(*D*), *off*′′′(*D*), *ch*′′′(*D*), *uch*′′′(*D*), *cat*′′′(*T*), *on*′′′(*T*), *off*′′′(*T*), *ch*′′′(*T*), *uch*′′′(*T*), and *cat*′′′(*D*) ∈ ℝ^n^ represent the corresponding parameter entries for the additional type 3 Michaelis–Menten processes. As before, the transition of intermediate complexes from the uncharged to the charged state requires activation; in the case of ATP production, this process is given by:

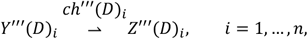

and in the case of ATP-hydrolysis

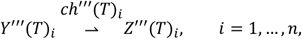

additional amounts of ATP, i.e. Δ*T*′*s*, are needed for the catalyzation of the respective processes. Δ*T* in the case of ATP-production is

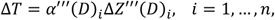

and in the case of ATP-hydrolysis

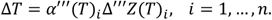

Again, 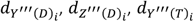, and 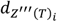, *i* = 1, …, *n*, represent the decay rates of the catalytic type 3 intermediate complexes formed between ribozymes and ADP or ATP, respectively.

##### The habitat restriction in the ATP-ADP cycle

The ATP demand rate *T*_*demand*_, representing the amount of ATP that must be hydrolyzed to ADP per unit of time to maintain the complete charging process in the system, is therefore given by:

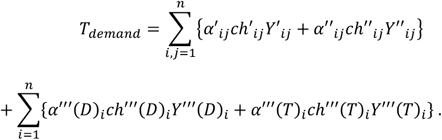

Conversely, the ATP–ADP cycle described by Eqs. (3), (4), and (5) determines the maximum total rate of ATP hydrolysis into ADP, which is given by:

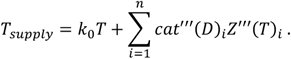

If *T*_demand_ > *T*_supply_, the resulting excess demand must be compensated for to restore equilibrium between ATP supply and consumption. To achieve this, all charging rates, 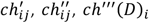, and *ch*′′′(*T*)_*i*_, are proportionally reduced by multiplication with a common damping factor *δ*:

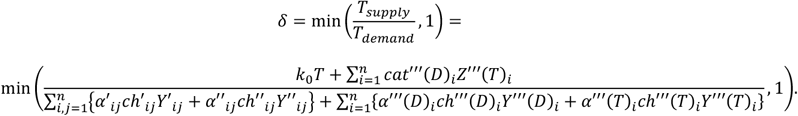

##### Allosteric ATP-regulation of ribozymes

In the following, we set *n* = 2 *m*, corresponding to an even number of ribozymes in the system. To incorporate allosteric ATP regulation into the model of ribozyme production, we assume that ATP molecules can bind to a non-active allosteric site on the ribozyme.

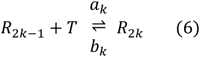

Here, *a*_*k*_ denotes the rate of allosteric ATP binding, and *b*_*k*_ the corresponding reverse rate, for *k* = 1, …, *m*. This distinction gives rise to two classes of ribozymes: ATP-loaded species—hereafter referred to as “loaded” (activated)—and ATP-unloaded species—hereafter referred to as “unloaded” (deactivated)—with respect to their allosteric sites. Accordingly, we assume that ribozymes *R*_*i*_with even indices *i* = 2*k* are allosterically ATP-loaded, whereas ribozymes with odd indices *i* = 2*k* − 1 remain unloaded. We further note that ribozymes *R*_2 *k-1*_ can interact with ATP in two distinct ways. Binding of ATP at the catalytic site results in the formation of a type 3 intermediate complex 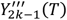, whereas binding at the allosteric site converts the ribozyme into its corresponding loaded form *R*_2*k*_.

An important property of the allosteric process described by Eq. (6) is that it is assumed to be non-catalytic. The transition between allosteric states is therefore not mediated by any of the three catalytic processes included in the model (types 1–3). In particular, type-1 catalysis does not modify the allosteric state of a ribozyme: ATP-loaded ribozymes cannot arise from ATP-free ribozymes, nor can ATP-free ribozymes be regenerated from ATP-loaded ribozymes, through type-1 catalytic reactions. Hence, type-1 catalysis does not contribute to the interconversion between ATP-loaded and ATP-free ribozyme populations. Based on this assumption, we define:

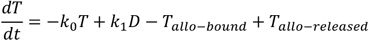

+ net release from intermediate complexes *T* with ribozymes *R*_*i*_

+ catalytic production of *T* from intermediate complexes with ribozymes *R*_*i*_ with *D*, and

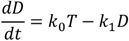

+ net release from intermediate complexes *D* with ribozymes *R*_*i*_

+ catalytic production of *D* from intermediate complexes with ribozymes *R*_*i*_ with *T*,

where

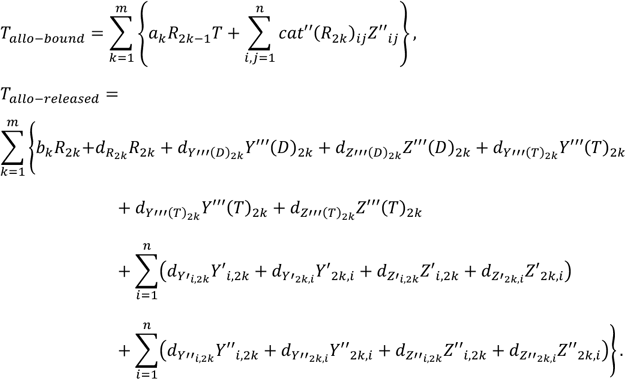

In the last equation, we assume that ATP bound at the allosteric site is released into the habitat during the decay of the ATP-loaded ribozyme *R*_2 *k*_, regardless of whether the ribozyme is present as a free species or as part of an intermediate complex.

Let finally *F*_*i*_, *i* = 1, … *n*, define the net conversion flow between the *R*_*2k-1*_′*s* and *R*_*2k*_′*s* according to Eq. (6):

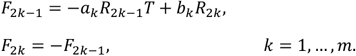

Now we can define the chemical dynamics of the whole system completely.

##### The explicit governing Michaelis-Menten equations

All these considerations lead to the following explicit ordinary Michaelis-Menten differential equations.

First, for the ATP-ADP cycle:

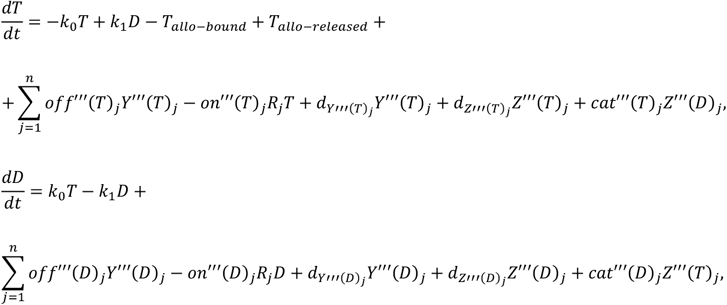

assuming that all ADP and ATP molecules bound to the catalytic intermediates are released into the habitat during the decay of their intermediate complexes, and

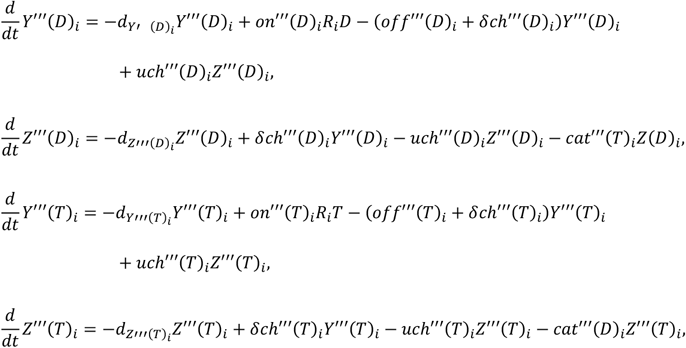

for *i* = 1, …, *n*.

Second, for the ribozymes:

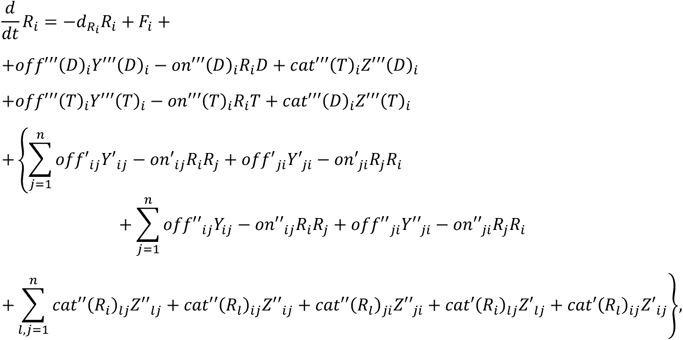

for *i* = 1, …, *n*;

and for the type 1 and 2 intermediate complexes, we have

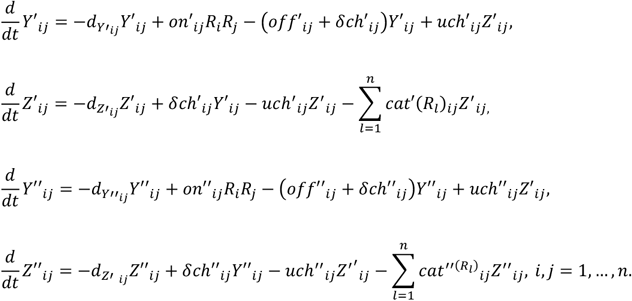

Finally, the total concentrations of the ribozymes, free plus bound, are given by

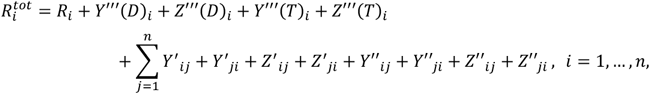

and, consequently, for the total amounts of ADP and ATP, free plus bound, it follows:

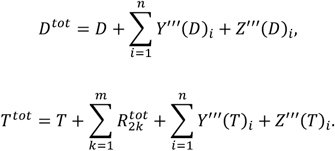

In the preceding analysis, we assumed that the external metabolism of the habitat constrains the total amount of these quantities to remain constant, such that:

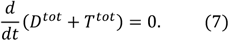

## 3. Results

We construct a minimal model of an autocatalytic ribozyme network capable of maintaining stable self-replication during the transition from geochemical prebiotic chemistry toward an emerging proto-biological system. The minimal requirements are (i) a prebiotically available source of activated chemical energy, represented here by an ATP/ADP-based energy-conversion cycle (Pinna et al., 2022), and (ii) autocatalytic, template-based self-sustained replication (Papastavrou et al., 2024; Gianni et al., 2026) that remains robust against the internal emergence of, or invasion by, parasitic ribozymes (Conrad et al., 2023; Pirovino et al., 2025).

Here, we introduce a specific realization of the general framework described in the Methods section (Eqs. (1)–(7)), which we refer to as the minimal stable model: an autocatalytic RNA-world habitat regulated by an ATP/ADP-based energy cycle. The construction of this model relies on the following assumptions. Because ribozyme replication is driven by an ATP/ADP-based proto-metabolic energy cycle (Pinna et al., 2022), the habitat must maintain a sufficient ATP supply to sustain replication. However, any prebiotic habitat has limited access to both metabolic energy and the molecular resources required for RNA synthesis. Since ATP serves simultaneously as an energy carrier and as a molecular resource for replication, we assume conservation of the total ATP and ADP pools, including both free and ribozyme-bound forms. Accordingly, the conservation relation defined by Eq. (7) also applies to the minimal model, as schematically illustrated in Fig. 1. As a starting point, we consider the shortest possible autocatalytic ribozyme cycle, namely a type-2 cycle of length one:

**Fig. 1.**
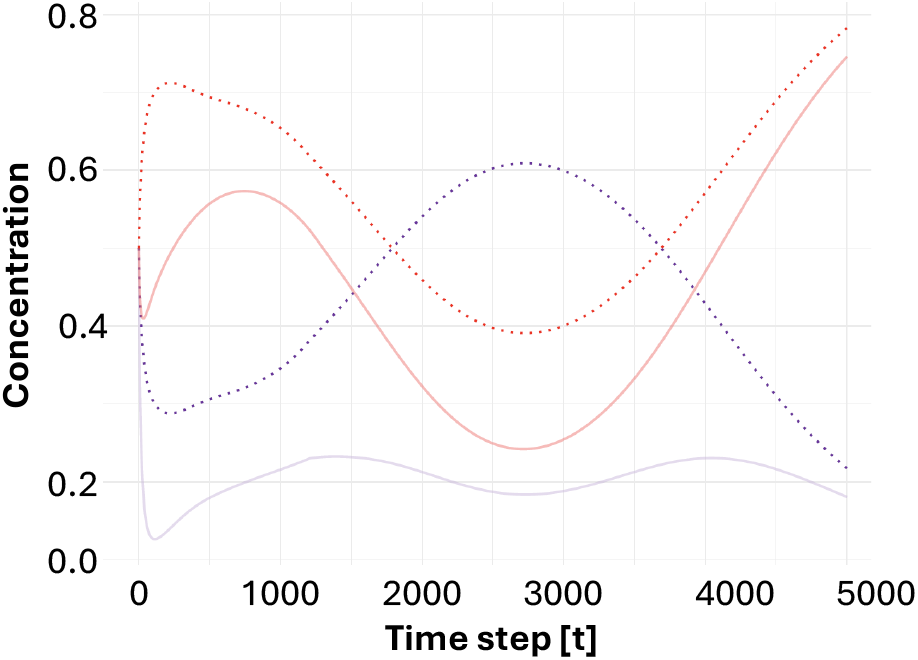
Minimal ATP–ADP cycle (schematic evolution). The habitat maintains a constant total nucleotide pool: *T* + *D* = *const*, with *T* = *T* + (*T*_tot_ − *T*) and *D* = *D* + (*D*_tot_ − *D*), with *T, D* denoting free and (*T* − *T*), (*D* − *D*) bound ATP and ADP, respectively. The y-axis shows the concentrations of ATP and ADP ribozymes, and the x-axis represents discrete time steps.

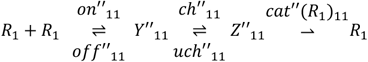

Here, *R*_1_ denotes the host replicase, which catalyzes its own replication. Throughout Figs. 2, we assume that replication of the host replicase is powered exclusively by an ATP/ADP-based chemical energy cycle naturally available within the habitat (Pinna et al., 2022). No ATP/ADP-cycle-catalyzing ribozyme is present at this stage. Consequently, free and total concentrations of ATP and ADP coincide, and the available ATP pool is directly coupled to replication of *R*_1_ (Figs. 2A–B).

**Figs. 2A–B.**
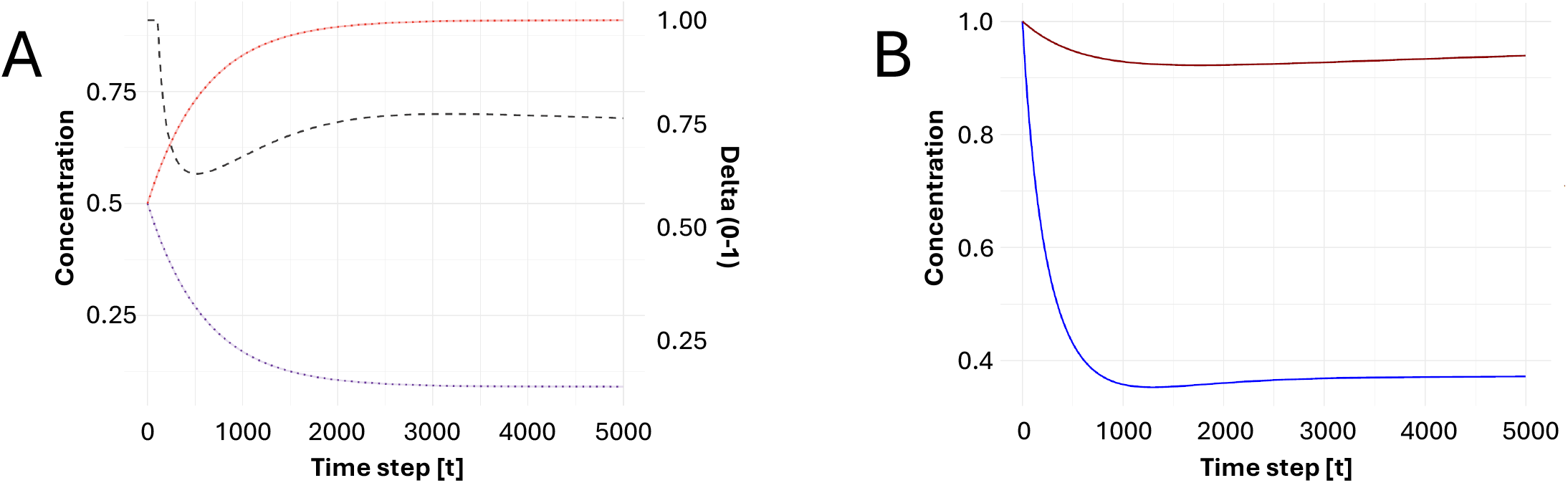
Prebiotic ATP production and *R*_1_replication. At moderate ATP consumption associated with replication, the system can be maintained without catalytic enhancement of the ATP–ADP cycle. (A) Free and total ATP and ADP concentrations are assumed to be identical. (B) All available ATP is immediately consumed by *R*_1_replication. (D_—), (D__tot_ − −), (delta_—), (T_—), (T__tot_− −); (*R*_1_, blue; total *R*_1,_, brown).

Despite its high standard free energy of hydrolysis of 50 kJ mol^-1^, ATP is chemically stable, with an estimated half-life of approximately one year under aqueous physiological-like conditions. In extant organisms, the ratio [ATP]/[ADP][Pi] is maintained approximately nine orders of magnitude away from equilibrium, reflecting the active regulation of cellular energy metabolism (Macdonald and Ashby, 2025). Under prebiotic conditions, however, such a strong thermodynamic disequilibrium would initially have been much less pronounced. In our model, the phosphorylation potential is subsequently increased through the evolution of enhanced ATP/ADP-cycle turnover, as described in the following section.

Because replication of the host replicase is inevitably error-prone, catalytically inactive parasitic ribozymes *R*_3_ continuously arise through mutation (Iranzo et al., 2016; Koonin et al., 2017, Shah et al., 2019). In the absence of an effective regulatory mechanism, uncontrolled replication of these parasites by the host replicase ultimately leads to collapse of the host–parasite system (*R*_1_, *R*_3_), (Figs. 3A-C). A major source of this instability is the characteristic binding behavior of parasites. Throughout this work, we define the parasitic binding property as a combination of a relatively high association rate with the host replicase *R*_1_, and very low association rates among parasitic ribozymes. As a result, parasitic ribozymes compete efficiently for access to the host replicase while exhibiting only weak self-association. This binding pattern gives rise to the characteristic evolutionary trajectory observed throughout our simulations: parasites initially proliferate at the expense of the host (Fig. 3C), but their continued expansion ultimately causes the collapse of both the host and parasite populations (Figs. 3B-C). Figs. 3 illustrate how the parasitic binding property, in the absence of any regulatory mechanism, inevitably drives the host–parasite system to collapse.

**Figs. 3A–C.**
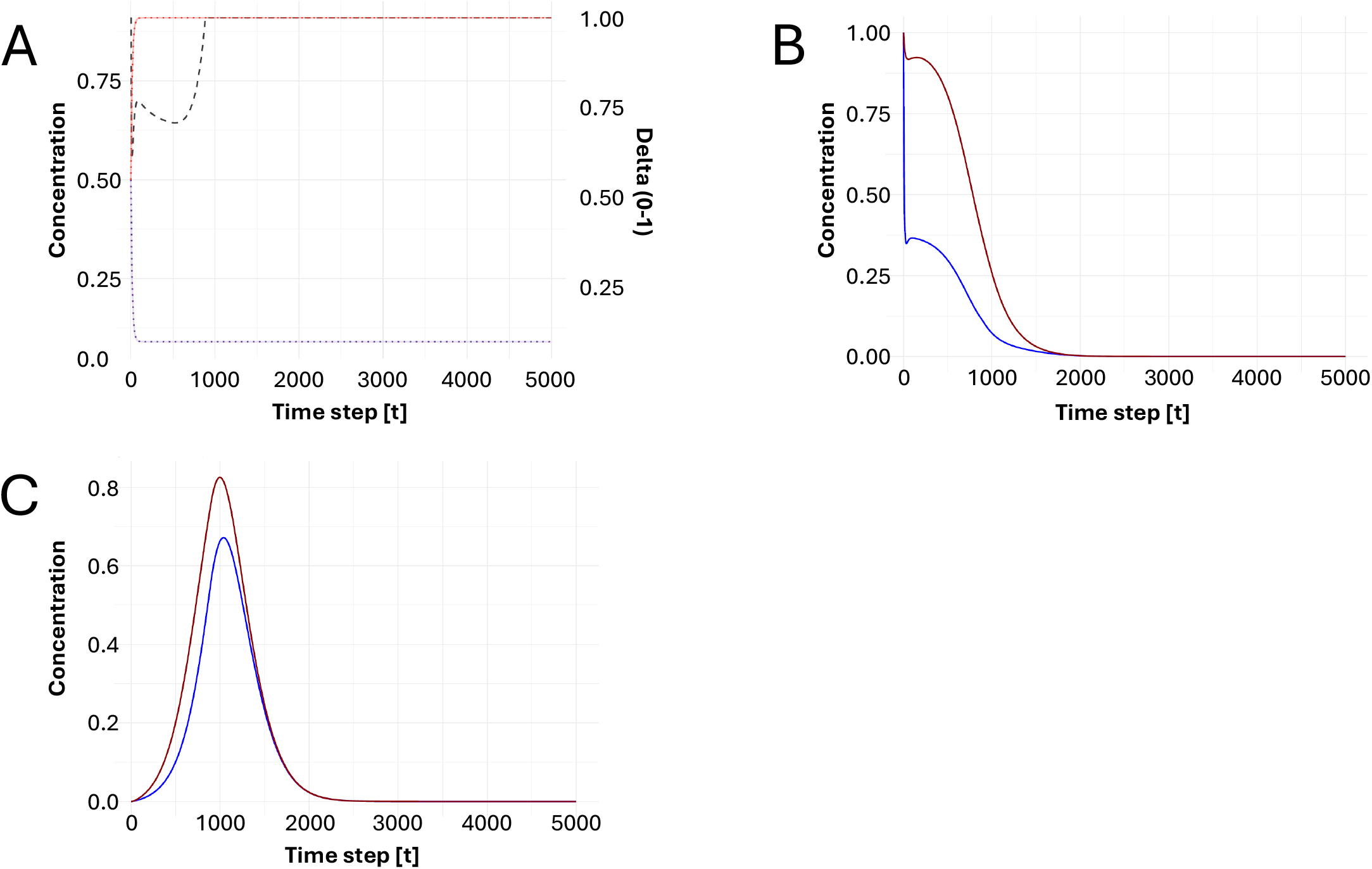
Catalytically inactive parasitic ribozyme *R*_3_. (A) Evolution of ADP (D_), ATP (T_), and (delta_); no ATP/ADP-cycle-catalyzing ribozyme is present at this stage. Initially, *R*_3_ proliferates at the expense of the host (C). However, unchecked parasite growth eventually causes the collapse of both the host *R*_1_population (B) and the *R*_3_parasite population (C). (D_—), (D__tot_ − −), (delta_—), (T_—), (T__tot_− −); (*R*_1_, blue; total *R*_1,_, brown; *R*_3_, blue; total *R*_3,_, brown).

We propose allosteric regulation as a plausible evolutionary mechanism by which early autocatalytic RNA systems could have mitigated molecular parasitism. This hypothesis is supported by the widespread occurrence of allosteric regulation in extant biological systems (Montserrat-Canals et al., 2025). According to Eq. (6), each ribozyme *R*_2 *k-1*_ may reversibly convert into an ATP-bound allosteric state, *R*_2 *k*_. The replicase *R*_1_ is assumed to lack an allosteric site and is therefore not subject to allosteric regulation (*R*_2_ = 0). This simplifying assumption isolates the effects of allosteric regulation on parasite control without introducing additional regulation of the replication process itself. Not all parasitic ribozymes are assumed to be allostery-prone. However, if a parasitic ribozyme *R*_3_undergoes ATP-induced conversion to the allosteric state *R*_4_, thereby completing the allosteric conversion cycle defined by Eq. (6), uncontrolled parasite proliferation can be markedly attenuated (Figs. 4A–D), provided that *R*_4_ possesses the assumed hyperparasitic binding property. In this scenario, allosteric regulation serves a dual evolutionary role. First, it suppresses runaway parasite proliferation. Second, rather than eliminating parasitic ribozymes, it creates an opportunity for their evolutionary exaptation into functional components of the molecular ecosystem, where they may ultimately contribute to host fitness.

**Figs. 4A-D.**
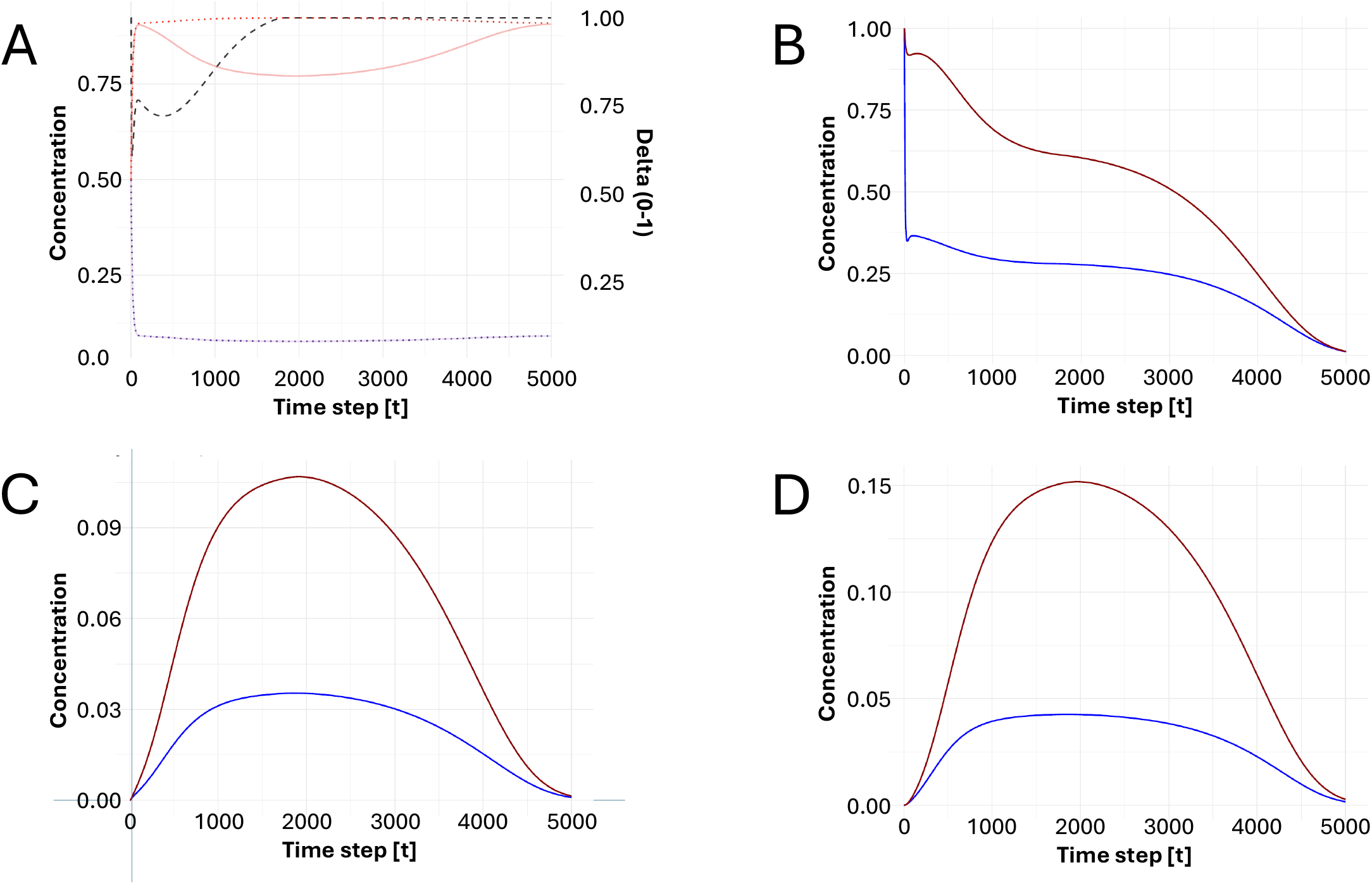
Allosteric regulation mitigates molecular parasitism. (A) Evolution of ADP (D_), ATP (T_), and (delta_); no ATP/ADP-cycle-catalyzing ribozyme is present at this stage. (B) Evolution of the host ribozyme *R*_1_. (C–D) The parasitic ribozyme *R*_3_undergoes ATP-induced allosteric conversion to *R*_4_, thereby substantially taming uncontrolled parasite proliferation, provided that *R*_4_satisfies the hyperparasitic binding property. (D_—), (D__tot_ − −), (delta_—), (T_—), (T__tot_− −); (*R*_1_, blue; total *R*_1,_, brown; *R*_3_, blue; total *R*_3,_, brown, *R*_4_, blue; total *R*_4,_, brown).

The hyperparasitic binding property is characterized by three binding preferences: a high association rate with the host replicase *R*_1_, comparable to that of the parasite *R*_3_; a high association rate with ATP-free parasitic ribozymes *R*_3_; and, optionally, an elevated association rate with ATP-bound parasitic ribozymes *R*_4_. Consequently, ATP-bound parasites not only compete with ATP-free parasites for access to the host replicase but also bind directly to parasitic ribozymes, thereby suppressing their uncontrolled proliferation. This hyperparasitic strategy, however, incurs a metabolic cost. ATP sequestration by the allosteric state *R*_4_reduces the concentration of free ATP, thereby slowing ATP–ADP turnover and progressively limiting the metabolic energy available to sustain RNA replication. Consequently, the metabolic energy available to sustain RNA replication progressively becomes limiting. When the ATP– ADP turnover rate is increased by doubling the rate constants *k*_0_and *k*_1_in Eq. (3) relative to the conditions shown in Figs. 4A–D, complete stabilization is restored (Figs. 5A–D). This finding indicates that the residual destabilization results primarily from reduced ATP turnover caused by ATP sequestration, rather than from allosteric regulation per se. The simulations therefore demonstrate that allosteric regulation can provide an effective mechanism for suppressing molecular parasitism, provided that ATP– ADP turnover remains sufficiently rapid to sustain the energetic demands of RNA replication.

**Figs. 5A-D.**
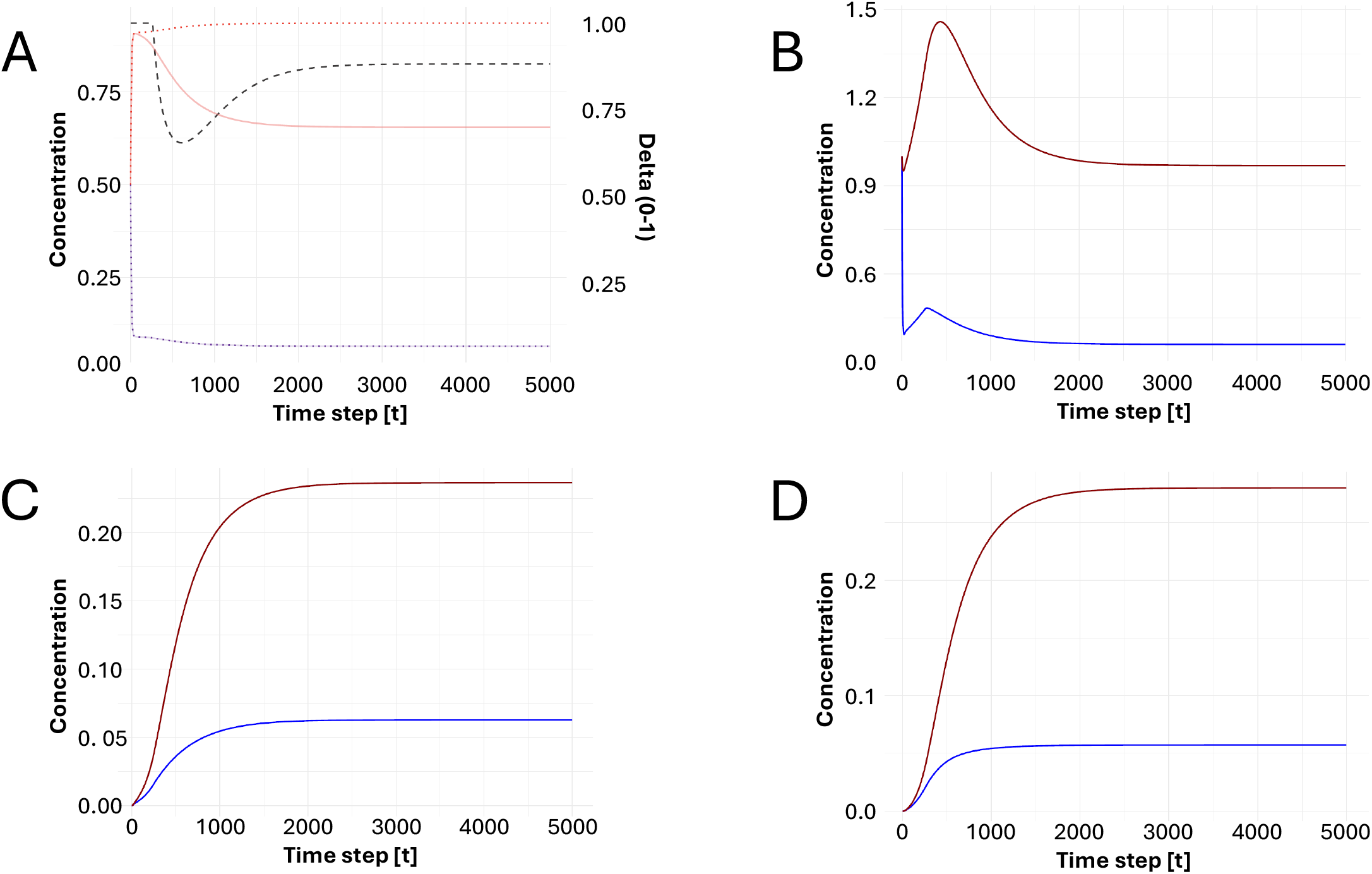
Accelerating the ATP/ADP cycle restores stabilization. (A) Evolution of ADP (D_), ATP (T_), and (delta_); no ATP/ADP-cycle-catalyzing ribozyme is present at this stage, but the ATP–ADP turnover rate is doubled. This increase, achieved by doubling the rate constants *k*_0_and *k*_1_, restores complete stabilization. (B–D) Evolution of the host ribozyme *R*_1_, parasitic ribozyme *R*_3_, and allosteric ribozyme *R*_4_, respectively. (D_—), (D__tot_ − −), (delta_—), (T_—), (T__tot_− −); (*R*_1_, blue; total *R*_1,_, brown; *R*_3_, blue; total *R*_3,_, brown, *R*_4_, blue; total *R*_4,_, brown).

In the first scenario, loss of replicase activity is intrinsically coupled to the acquisition of an allosteric site. Consequently, every parasitic product of the host replicase *R*_1_ immediately enters the allosteric conversion cycle described by Eq.(6). Figs. 4A-D and 5A-D demonstrate that this mechanism effectively stabilizes the host–parasite system. If such an intrinsic coupling were readily selectable in an RNA-world habitat, it would constitute the most energy-efficient solution to molecular parasitism.

The second scenario is energetically less efficient but is likewise supported by our simulations. In this scenario, a parasitic ribozyme, denoted *R*_5_, evolves into an allosterase (Fig. 6D). For simplicity, we assume that the allosterase is itself allostery-prone and therefore possesses an ATP-bound allosteric counterpart, *R*_6_(Fig. 6E). Upon encounter, *R*_5_, and possibly also *R*_6_, catalyzes the conversion of a non-allostery-prone parasitic ribozyme *R*_3_into the allostery-prone forms *R*_3_and *R*_8_through a type-1 catalytic process (Figs. 6F–G). Figures 6A–G show that this mechanism substantially improves system stability relative to the unregulated dynamics shown in Figs. 3A–C. Complete stabilization, however, is not achieved because ATP sequestration during allosteric regulation slows ATP–ADP turnover, thereby limiting the metabolic energy available for RNA replication. When the ATP–ADP turnover rate is increased by tripling the rate constants *k*_0_and *k*_1_, complete stabilization is restored (Figs. 7A–G).

**Figs. 6A-G.**
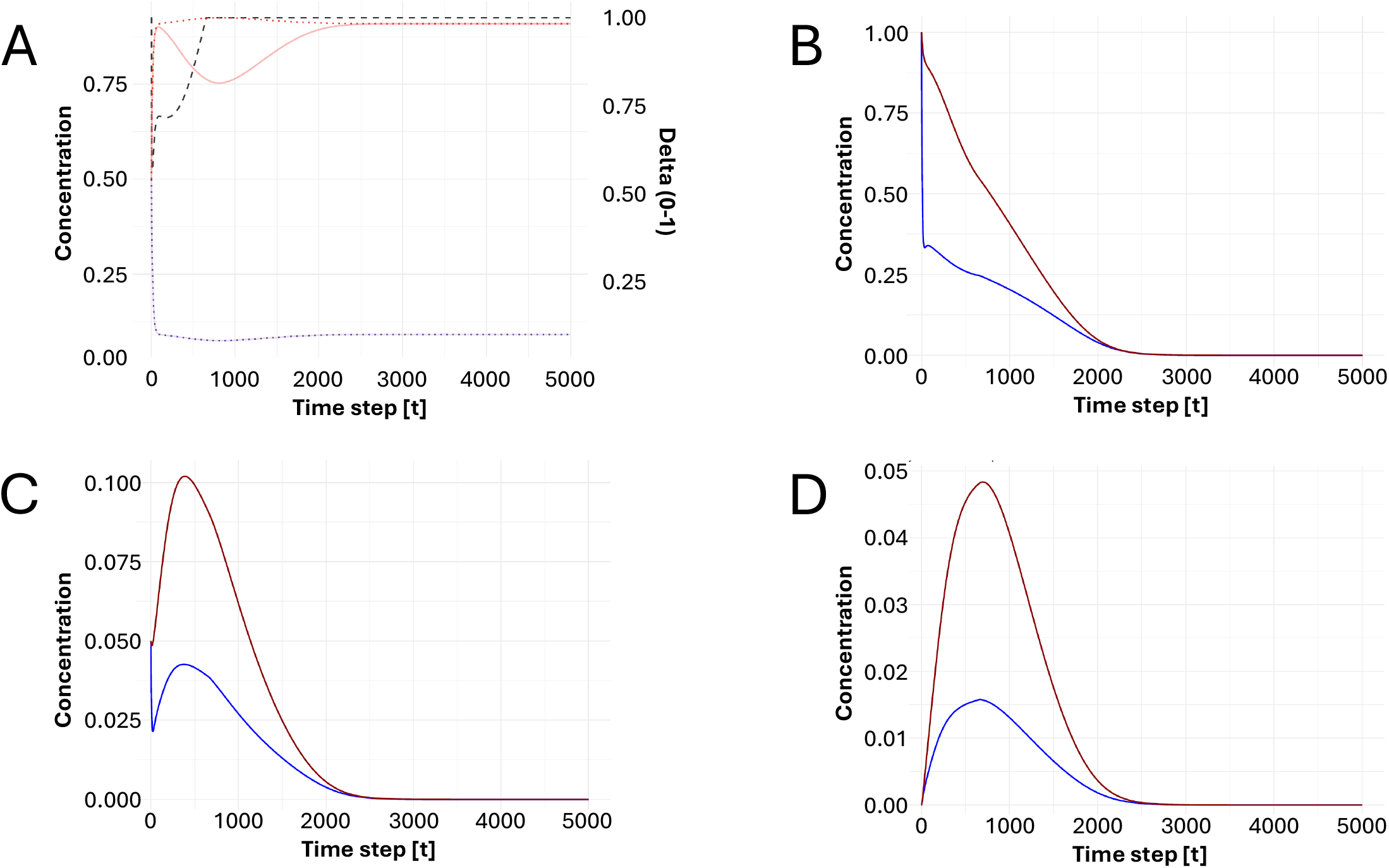
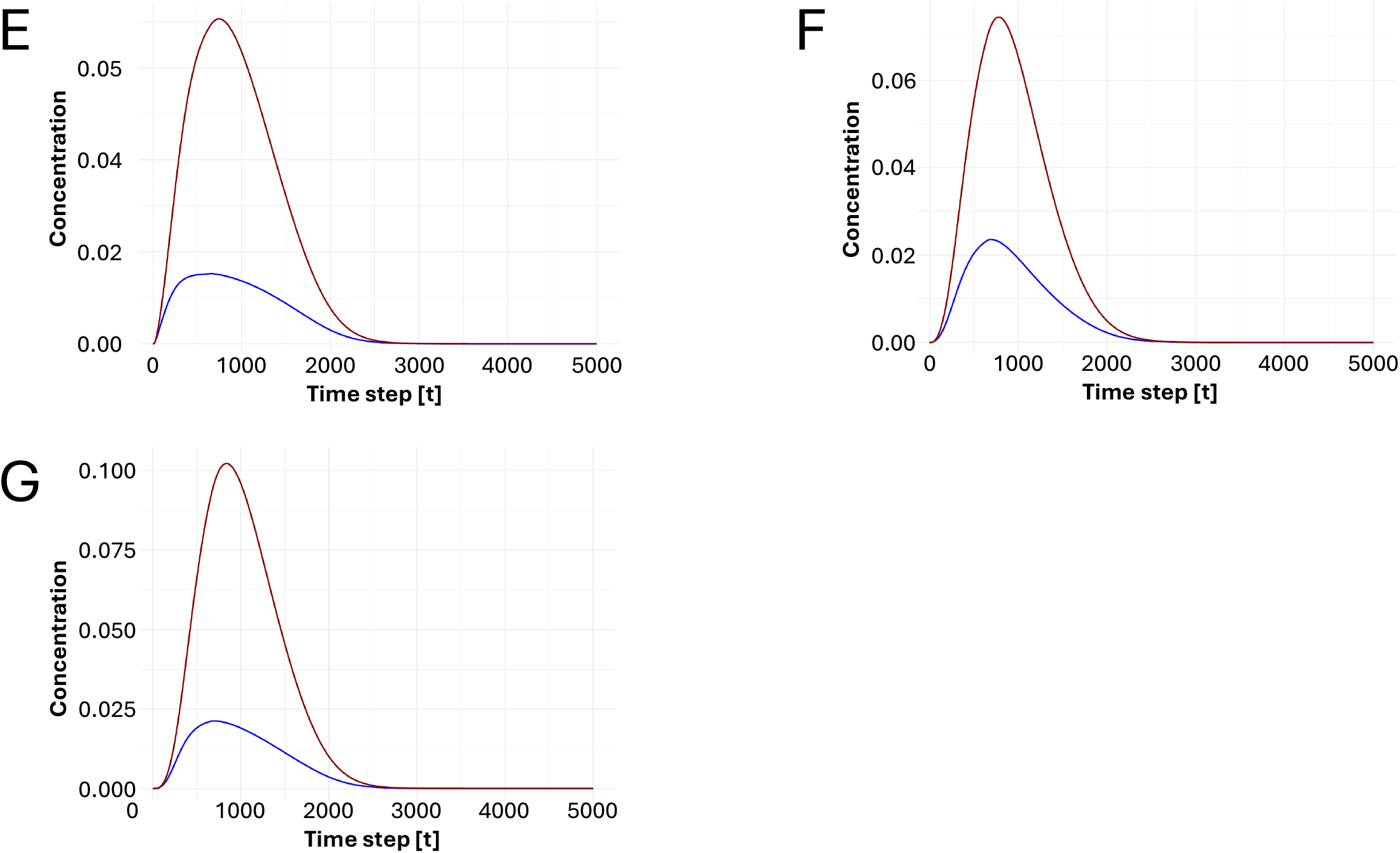
Allosterases confer allostery in trans. (A) Evolution of ADP (D_), ATP (T_), and (delta_), no ATP/ADP-cycle-catalyzing ribozyme is present at this stage. The allosterase *R*_5_undergoes ATP-induced conversion to its allosteric counterpart *R*_6_, which subsequently converts parasitic ribozymes *R*_3_into the allostery-prone forms *R*_3_and *R*_8_. This mechanism improves system stability but does not achieve complete stabilization. (B–G) Evolution of *R*_1_, *R*_3_, *R*_S_, *R*_6_, *R*_?_, and *R*_8_, respectively. (D_—), (D__tot_ − −), (delta_—), (T_—), (T__tot_− −); (*R*_1_, blue; total *R*_1,_, brown; *R*_3_, blue; total *R*_3,_, brown; *R*_S_, blue; total *R*_5,_, brown; *R*_6_, blue; total *R*_6,_, brown; *R*_3_, blue; total *R*_3,_, brown; *R*_8_, blue; total *R*_8,_, brown).

**Figs. 7A-G.**
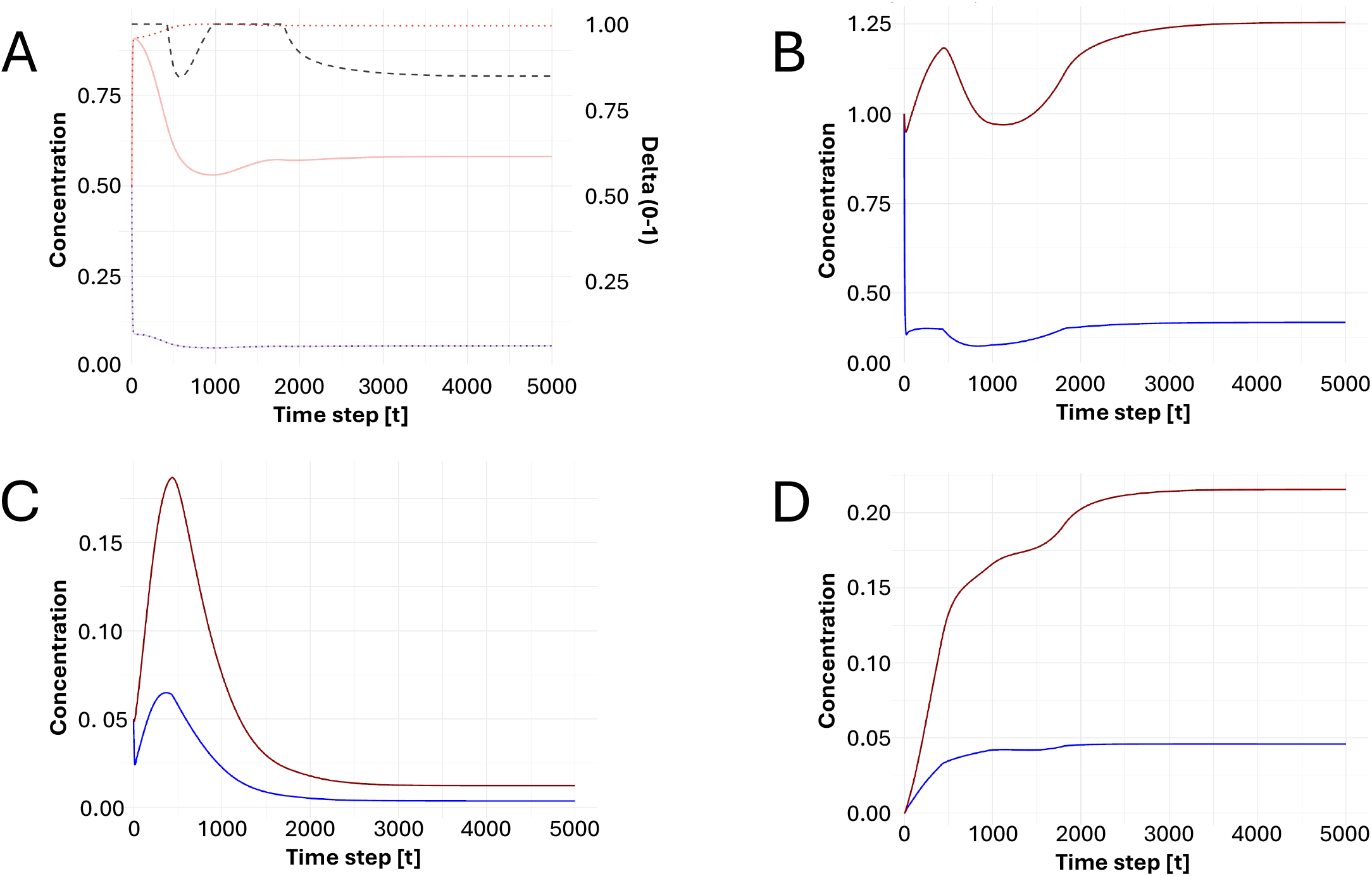
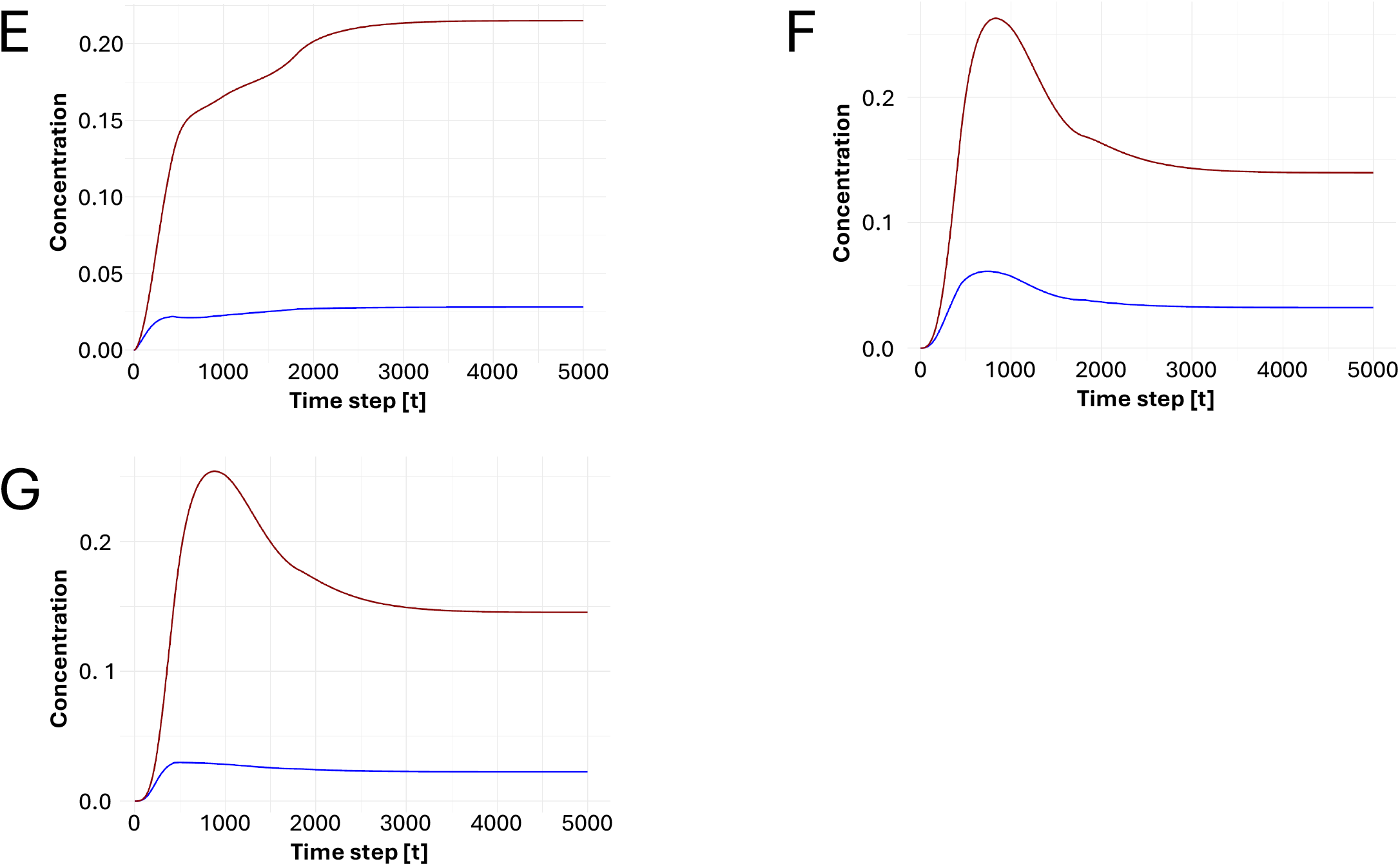
Allosterases and enhanced ATP–ADP turnover. (A) Evolution of ADP (D_), ATP (T_), and (delta_), no ATP/ADP-cycle-catalyzing ribozyme is present at this stage, but the turnover rate of the ATP/ADP cycle is tripled. The allosterase *R*_5_undergoes ATP-induced conversion to its allosteric counterpart *R*_6_, which subsequently catalyzes the conversion of parasitic ribozymes *R*_3_into the allostery-prone forms *R*_3_and *R*_8_. Under these conditions, complete stabilization is achieved. (B–G) Evolution of *R*_1_, *R*_3_, *R*_S_, *R*_6_, *R*_?_, and *R*_8_, respectively. (D_—), (D__tot_ − −), (delta_—), (T_—), (T__tot_− −); (*R*_1_, blue; total *R*_1,_, brown; *R*_3_, blue; total *R*_3,_, brown; *R*_5_, blue; total *R*_5,_, brown; *R*_6_, blue; total *R*_6,_, brown; *R*_3_, blue; total *R*_3,_, brown; *R*_8_, blue; total *R*_8,_, brown).

Taken together, these simulations demonstrate that two distinct evolutionary pathways can stabilize RNA replication in the presence of molecular parasitism. The first relies on intrinsic allostery-proneness, in which all newly emerging parasitic ribozymes are inherently capable of ATP-dependent allosteric regulation (Figs. 4 and 5). The second relies on allosterase-mediated regulation, in which a specialized allosterase converts non-allostery-prone parasitic ribozymes into allostery-prone forms (Figs. 6 and 7). In both scenarios, molecular parasitism is tamed through ATP-dependent allosteric regulation. However, ATP sequestration by the allosteric states reduces the effective ATP–ADP turnover rate, thereby imposing an energetic cost on the system.

The simulations presented above show that allosteric regulation effectively tames molecular parasitism but simultaneously reduces the metabolic energy supply by sequestering ATP. This trade-off creates a selective pressure favoring ribozymes that restore ATP–ADP turnover. We therefore propose that, among the predominantly catalytically inactive parasitic ribozymes, one mutant *R*_3_acquires catalytic activity for the ATP–ADP cycle (type 3), thereby functioning as an ATP synthase/ATPase (Figs. 8A– D).

**Figs. 8A-D.**
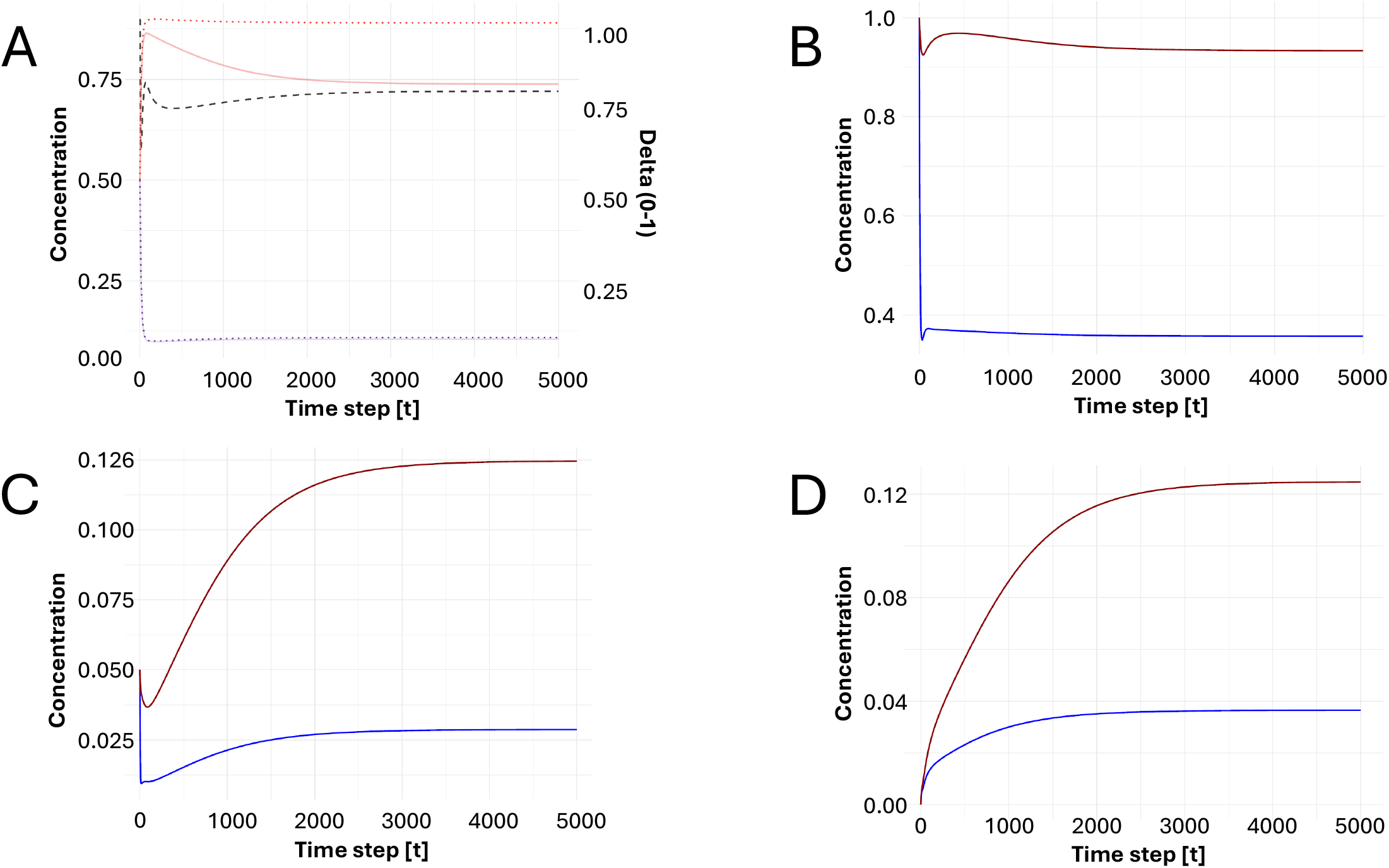
Parasite ribozymes *R*_3_evolve into ATPase/ATP synthetase ribozymes. (A) Evolution of ADP (D_), ATP (T_), and (delta_); ATPase/ATP synthase activity has emerged from the parasitic ribozyme *R*_3_(C). As shown in Fig. 4, ATP-dependent allosteric regulation by *R*_4_(D) tames parasitism by *R*_3_, but requires additional ATP turnover that is supplied by the newly evolved ATPase/ATP synthase activity. (B– D) Evolution of *R*_1_, *R*_3_, and *R*_4_, respectively. (D_—), (D__tot_ − −), (delta_—), (T_—), (T__tot_− −); (*R*_1_, blue; total *R*_1,_, brown; *R*_3_, blue; total *R*_3,_, brown, *R*_4_, blue; total *R*_4,_, brown).

Following the first evolutionary scenario described in the previous section, we assume that this ATP synthase/ATPase ribozyme is itself allostery-prone and therefore possesses the ATP-bound allosteric counterpart *R*_4_. As shown in Figs. 4 and 5, allosteric regulation alone substantially improves the stability of the host–parasite system but does not achieve complete stabilization because ATP sequestration slows ATP–ADP turnover. By simultaneously catalyzing ATP regeneration, *R*_3_compensates for this reduction in ATP turnover, thereby restoring the metabolic energy supply required for RNA replication and enabling complete stabilization of the system.

The allosteric regulation of the ATP synthase/ATPase ribozyme *R*_3_ serves two complementary functions, namely first, control of molecular parasitism. The ATP-loaded allosteric counterpart *R*_4_ attenuates the parasitic behavior of *R*_3_ through the hyperparasitic binding property. Second, regulation of ATP availability. Because *R*_4_ is catalytically inactive, ATP bound to its allosteric site is temporarily removed from the metabolically active pool. The allosteric counterpart therefore acts as a dynamic ATP buffer, regulating ATP availability while *R*_3_ maintains ATP–ADP turnover. Together, these two functions establish a feedback mechanism in which ATP synthase/ATPase activity maintains the metabolic energy supply required for RNA replication, whereas allosteric regulation prevents uncontrolled proliferation of parasitic ribozymes.

Conceptually, our model suggests the following possible evolutionary sequence: (i) increased energy throughput via the emergence of more efficient mechanisms of energy capture and conversion; (ii) stabilization of autocatalytic replication networks against exploitation by parasitic or non-productive molecular variants (Eigen and Schuster, 1979; Maynard Smith and Szathmáry, 1995); (iii) optimization of catalytic efficiency only after regulatory mechanisms have emerged that ensure that increased catalytic activity primarily benefits the productive network (Buss, 1987; Nowak, 2006). This sequence reflects a control-before-optimization principle emerging from our model: in evolving systems containing exploitable interactions, increases in efficiency may initially amplify the success of parasitic variants rather than enhance the stability of the system as a whole (Szathmáry, 2015). Once mechanisms evolve that suppress exploitation and align the fitness interests of molecular components with persistence of the higher-level network, further increases in catalytic efficiency become selectively advantageous. A similar logic is found in theories of evolutionary transitions in individuality, where the emergence of higher levels of biological organization requires mechanisms that constrain the proliferation of selfish elements and stabilize cooperation (Maynard Smith and Szathmáry, 1995; Szathmáry, 2015). More generally, this concept parallels principles from control theory and systems science, in which regulation, feedback, and error control provide the foundation upon which performance optimization can subsequently occur (Wiener, 1948; Krakauer et al., 2020).

Once an allostery-prone ATP synthase/ATPase ribozyme *R*_3_has evolved and maintains a sufficiently high ATP–ADP turnover rate, the host–parasite system remains stable despite the subsequent emergence of additional parasitic ribozymes. Figures 9A–F show that stabilization is maintained following the appearance of further parasitic mutants *R*_5_and *R*_6_, provided that these newly arising parasites are likewise allostery-prone. These simulations indicate that stabilization is not restricted to the initial host–parasite interaction but extends to successive rounds of parasite emergence. ATP-dependent allosteric regulation therefore constitutes a general mechanism for maintaining the long-term stability of an autocatalytic RNA network in the face of recurrent mutational invasion.

**Figs. 9A-F.**
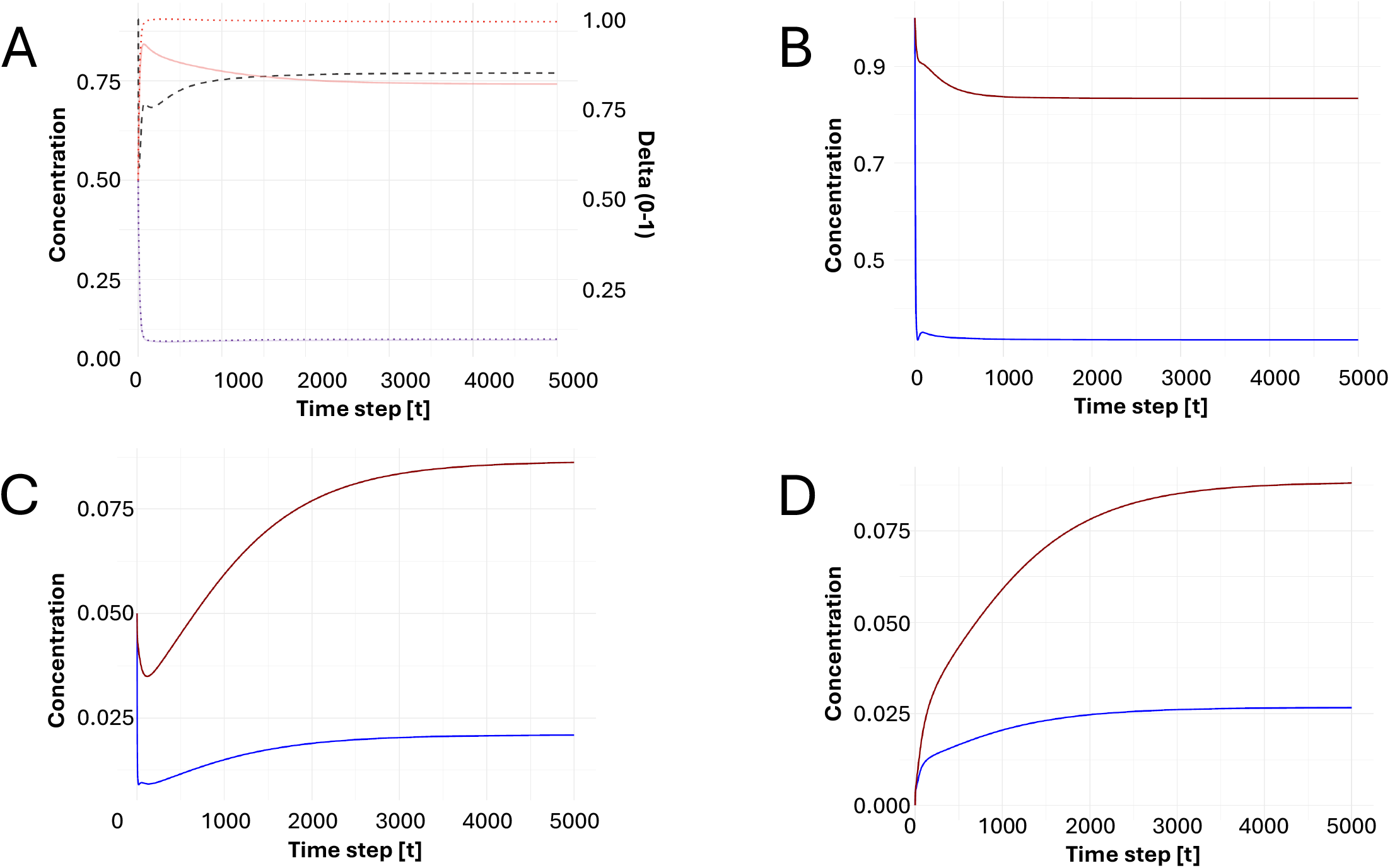
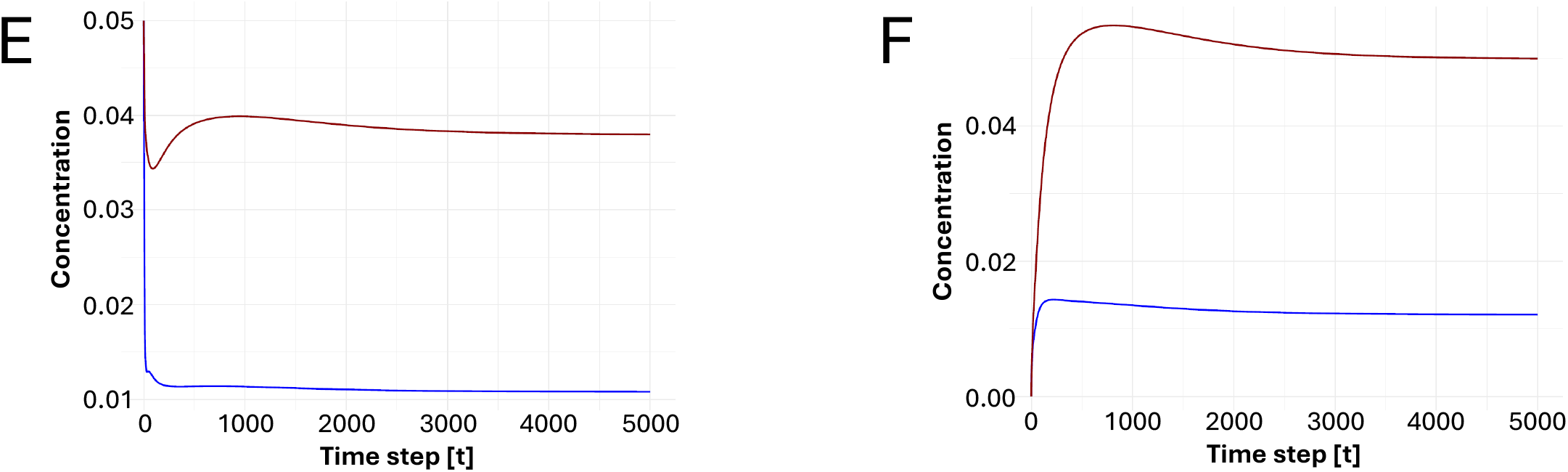
ATPase/ATP synthase ribozymes enable resistance to subsequent parasite invasion. (A) Evolution of ADP (D_), ATP (T_), and (delta_); ATPase/ATP synthase activity has emerged from the parasitic ribozyme *R*_3_(C). The resulting allosterically regulated habitat (D), supported by sufficient ATP supply (A), resists subsequent invasion by parasitic ribozymes *R*_5_and *R*_6_(E–F), provided that newly emerging parasites are allostery-prone. (B–F) Evolution of *R*_1_, *R*_3_, *R*_4_, *R*_5_, and *R*_6_, respectively. (D_—), (D__tot_ − −), (delta_—), (T_—), (T__tot_− −); (*R*_1_, blue; total *R*_1,_, brown; *R*_3_, blue; total *R*_3,_, brown, *R*_4_, blue; total *R*_4,_, brown, *R*_5_, blue; total *R*_5,_, brown, *R*_6_, blue; total *R*_6,_, brown).

The previous section identified two evolutionary mechanisms by which molecular parasitism can be controlled in simple host–parasite systems: newly emerging parasitic ribozymes are intrinsically allostery-prone; or non-allostery-prone parasitic ribozymes are converted into allostery-prone forms by a specialized allosterase. Subsequently, it was demonstrated that the first mechanism remains effective in the presence of successive waves of parasitic mutation. Here we show that the same conclusion holds for the second mechanism. Specifically, the combined system consisting of the host replicase *R*_1_, the ATP synthase/ATPase pair *R*_3_, *R*_4_ and the allosterase pair *R*_5_, *R*_6_ remains stable following the emergence of the additional non-allostery-prone parasitic ribozyme *R*_3_, which is subsequently converted into allostery-prone forms *R*_9_, *R*_10_ (Figs. 10A-I). The simulations therefore demonstrate that allosterase-mediated regulation provides a robust stabilization mechanism not only for simple host–parasite systems but also for progressively more complex autocatalytic RNA networks. This completes our analysis of the mathematical simulations of the minimal model derived from the general framework defined by Eqs. (1)–(7).

**Figs. 10A-I.**
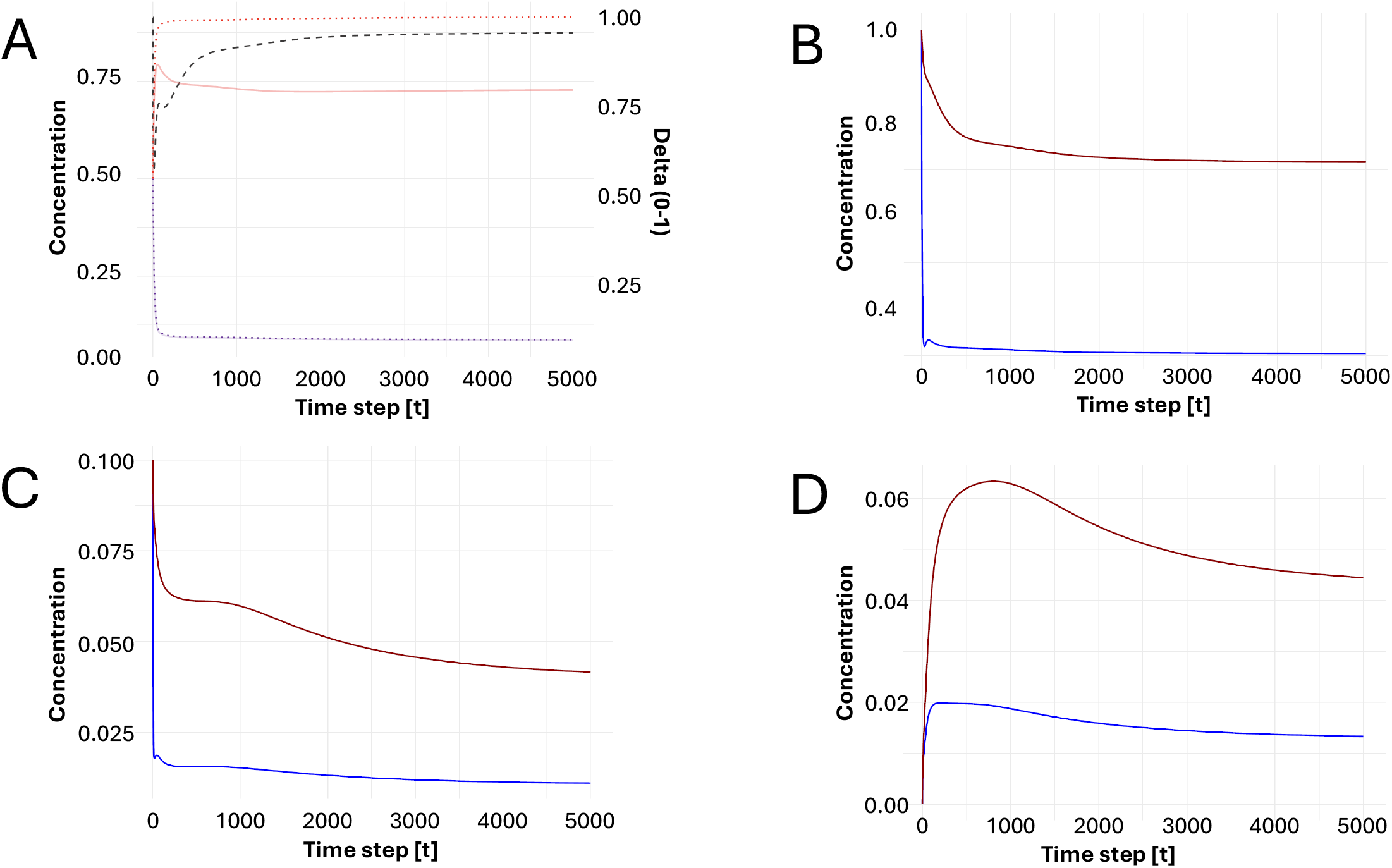
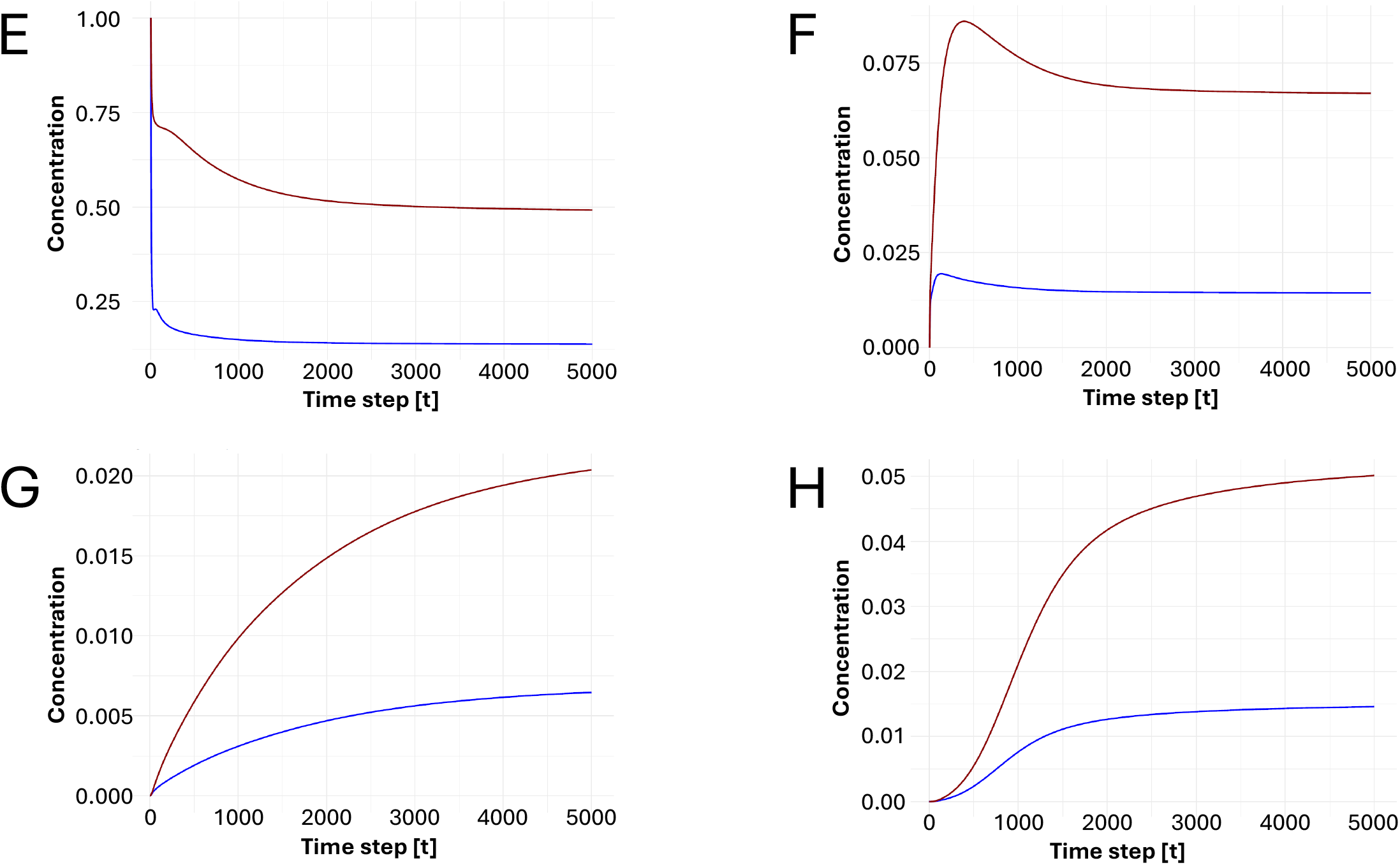
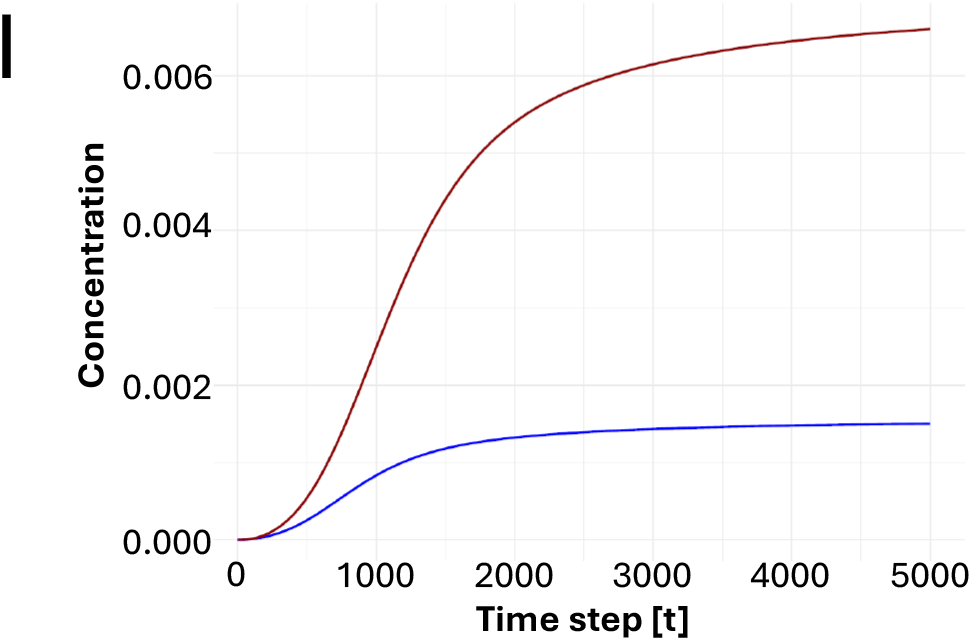
Habitat with ATPase/ATP synthetase ribozymes and allosterases recapitulates parasite resistance. (A) Evolution of ADP (D_), ATP (T_), and (delta_); ATPase/ATP synthase activity emerges from the parasitic ribozyme *R*_3_(C). The resulting integrated system, comprising the host replicase *R*_1_(B), the ATP synthase/ATPase ribozyme *R*_3_and its allosteric counterpart *R*_4_(C–D), and the allosterase pair *R*_5_and *R*_6_(E–F), remains stable following the emergence of the non-allostery-prone parasitic ribozyme *R*_3_(G). The allosterase subsequently converts *R*_3_into the allostery-prone forms *R*_9_and *R*_10_ (H–I), thereby maintaining parasite resistance. (D_—), (D__tot_ − −), (delta_—), (T_—), (T__tot_− −); (*R*_1_, blue; total *R*_1,_, brown; *R*_3_, blue; total *R*_3,_, brown, *R*_4_, blue; total *R*_4,_, brown, *R*_5_, blue; total *R*_5,_, brown, *R*_6_, blue; total *R*_6,_, brown, *R*_3_, blue; total *R*_3,_, brown, *R*_9_, blue; total *R*_9,_, brown, *R*_10_, blue; total *R*_10,_, brown).

The aim of this work was to identify the smallest autocatalytic RNA network capable of sustained self-replication while remaining robust against the inevitable emergence of non-catalytic parasitic mutants. Our simulations show that such a system requires four essential components: a host replicase *R*_1_ capable of self-replication; an ATP synthase/ATPase ribozyme *R*_3_ that maintains ATP–ADP turnover by catalyzing the metabolic energy cycle; ATP-dependent allosteric regulation of parasitic ribozymes through the hyperparasitic binding property; and a mechanism that ensures all parasitic ribozymes become allostery-prone, either intrinsically through an evolutionarily accessible molecular mechanism, or through the emergence of an allosterase that converts non-allostery-prone parasites into allostery-prone forms.

Accordingly, under the first evolutionary scenario, the minimal model consists of the host replicase *R*_1_, the ATP synthase/ATPase pair *R*_3_, *R*_4_, and additional allostery-prone parasitic ribozymes. In the second evolutionary scenario, the minimal model additionally contains an allosterase pair *R*_5_, *R*_6_, which converts newly emerging non-allostery-prone parasitic ribozymes into allostery-prone forms. The minimal model can, of course, be extended in several directions. For example, the replication cycle may require multiple ribozymes, or additional type-1 to type-3 catalytic interactions may evolve among catalytically active and catalytically inactive parasitic ribozymes.

Previous work (Pirovino et al., 2025) demonstrated that autocatalytic RNA networks can be stabilized when the replicase generates sufficient numbers of hyperparasitic ribozymes with binding properties similar to those considered here, assuming that the metabolic energy supply is provided entirely by the surrounding habitat. The present study extends this concept in two fundamental respects. First, we introduce an ATP-dependent allosteric regulatory mechanism that couples the metabolic state of the habitat directly to the replication dynamics of the RNA network, rather than treating metabolism as an externally maintained process. Whereas earlier work (Conrad et al., 2023) examined parasite-mediated regulation of metabolic energy supply using microbiological Lotka–Volterra-type models, we formulate the problem here within a thermodynamically consistent chemical framework based on Michaelis–Menten kinetics. Second, we identify a minimal autocatalytic RNA system in which ATP-dependent allosteric regulation fulfills a dual role: it maintains the metabolic energy supply required for RNA replication while simultaneously enabling the emergence of hyperparasitic regulation that tames the inevitable proliferation of catalytically inactive parasitic ribozymes.

More broadly, the present model suggests that the emergence of a stable RNA-based proto-biological system requires the integration of three fundamental properties: information replication, energetic autonomy, and internal regulation. Self-replication alone is insufficient, because inevitable mutational errors generate parasitic molecular variants that can exploit the replication machinery. Conversely, energy acquisition alone does not guarantee persistence, because increased energetic throughput can accelerate both productive and parasitic processes. Stability therefore emerges only when energy conversion becomes coupled to regulatory mechanisms that control molecular conflicts within the replicating network.

In this framework, allosteric regulation represents a possible early form of molecular governance: a mechanism by which the replication system begins to regulate access to limited energetic resources and constrain the proliferation of non-productive components. The emergence of ATP/ADP-cycle-catalyzing ribozymes and allosteric control therefore establishes a feedback architecture in which metabolic activity, replication dynamics, and parasite suppression become mutually coupled.

The minimal model should not be interpreted as a unique historical pathway for the origin of life, but rather as a demonstration of a possible route by which simple autocatalytic RNA systems could overcome fundamental stability constraints. More elaborate prebiotic systems may have involved multiple replicases, alternative energy currencies, compartmentalization, or additional catalytic interactions. Nevertheless, the principles identified here—energy supply, conflict regulation, and selective stabilization of productive interactions—represent general requirements for the transition from molecular replication toward autonomous evolving systems.

## 4. Discussion

Origin-of-life models have traditionally emphasized either replication-first or metabolism-first scenarios. In contrast, the present work investigates how replication, energy conversion, and molecular regulation may have co-evolved as mutually dependent processes within a single autocatalytic RNA network. We formulate this problem within a Michaelis–Menten framework in which ATP/ADP-based energy conversion drives template-directed ribozyme replication, while ATP-dependent allosteric regulation provides dynamic control over molecular interactions. Rather than treating metabolism as an externally supplied resource, the model explicitly couples ATP turnover to the replication dynamics of the autocatalytic network itself.

The central result is the identification of a minimal stable architecture capable of sustained self-replication despite the continual emergence of non-catalytic parasitic mutants. This architecture requires a self-replicating host ribozyme, a ribozyme that catalyzes ATP/ADP turnover, and ATP-dependent allosteric regulation acting on parasitic ribozymes through the hyperparasitic binding property. Together, these elements establish a feedback loop in which energy conversion, replication, and parasite suppression become dynamically integrated.

An important outcome of the model is the identification of an intrinsic energetic trade-off. Allosteric regulation suppresses parasite proliferation but simultaneously sequesters ATP, thereby reducing ATP/ADP turnover and limiting the energy available for replication. Stable autocatalytic growth therefore requires compensation through increased ATP regeneration. In our simulations, this compensation is provided by the evolution of an ATP/ADP-cycle-catalyzing ribozyme. The resulting coupling between regulation and energy conversion illustrates that metabolic innovation and regulatory innovation are not independent processes but may have evolved in response to one another.

The model further suggests a possible evolutionary sequence in which increased energetic throughput is followed by the emergence of regulatory mechanisms that suppress molecular parasitism, thereby permitting subsequent optimization of catalytic efficiency. This control-before-optimization principle is consistent with broader theories of evolutionary transitions, in which higher levels of biological organization become possible only after mechanisms evolve that limit internal conflict and stabilize cooperation (Maynard Smith and Szathmáry, 1995; Szathmáry, 2015). Although the present model does not establish the historical order of these events, it demonstrates that such a sequence provides a plausible route toward stable RNA-based evolution.

The simulations also identify two feasible mechanisms by which allosteric control may have emerged. In one scenario, parasitic ribozymes intrinsically acquire allosteric regulation following loss of replicase activity. In the other, an allosterase ribozyme evolves that catalyzes the conversion of non-allostery-prone parasites into allostery-prone forms. Both mechanisms produce comparable stabilization of the replication network, suggesting that robust parasite control does not depend on a unique evolutionary solution but may arise through alternative regulatory pathways.

To our knowledge, this is among the first mathematical models to integrate template-directed RNA replication, ATP/ADP-based energy conversion, and ATP-dependent allosteric regulation within a single autocatalytic reaction network. The model should not be interpreted as a reconstruction of the unique historical origin of life, but rather as a demonstration that stable autocatalytic evolution can emerge from the dynamic coupling of information replication, proto-metabolism, and molecular regulation. Importantly, this finding proved robust across the exploration of alternative model formulations. Scenarios in which catalytic performance was enhanced without the emergence of regulatory control—including an initially more efficient ATP/ADP cycle—consistently failed to generate a stable, self-maintaining habitat. By contrast, the appearance of ATP-dependent regulatory interactions repeatedly enabled the establishment of persistent autocatalytic evolution. These results therefore suggest that regulatory innovation was not simply an accessory refinement of an already functional metabolic network, but a prerequisite for the transition from transient chemical self-organization to autonomous, evolvable proto-biological systems. Within the limits of this modeling framework, they support the view that the emergence of regulation constituted a critical systems-level transition, allowing metabolic activity to become sufficiently coordinated and resilient to sustain long-term evolutionary dynamics.

## Declaration of generative AI and AI-assisted technologies in the manuscript preparation process

During the preparation of this work, the first author used Chat-GPT (OpenAI) to improve language clarity and structure. After using this tool, the author(s) reviewed and edited the content as needed and take(s) full responsibility for the content of the published article.

## CRediT authorship contribution statement

Bernard Conrad: Writing – original draft, review & editing, Methodology, Investigation, Supervision, Conceptualization. Christian Iseli: Writing – review & editing, Software, Formal analysis. Magnus Pirovino: Writing – original draft, review & editing, Formal analysis, Methodology, Software, Conceptualization.

## Data and code availability

The R code implementing the equations and reproducing the analyses presented in this study is available on https://github.com/BICC-UNIL-EPFL/energy-metabolism-and-hyperparasitism and is archived at https://doi.org/10.5281/zenodo.20408570.

## Acknowledgements

We warmly thank Udo Seifert for helpful discussions and his insights into statistical thermodynamics.

## Appendix

### Parametrizations and initial values for model calculations

#### 1.1 Tables of relevant parameters for the respective Michaelis-Menten processes

The model parameters and initial conditions are defined as dimensionless quantities because the objective is to investigate the qualitative and quantitative behavior of the system rather than to calibrate the model to a particular biochemical setting. Nevertheless, the relative parameter values are chosen to be biologically plausible. For example, in Eq. (3), describing the intrinsic ATP–ADP cycle, the ratio of the transformation rates *k*_0_ and *k*_1_is selected so that the steady-state ATP-to-ADP ratio falls between 2 and 10, in agreement with values commonly observed in living cells (Hardie, D.G., 2003).

The terminology used for the different classes of kinetic parameters—namely association (on) rates, dissociation (off) rates, charging (activation) rates, discharging/uncharging (deactivation) rates, and catalytic rates—is consistent with that introduced in the Methods section. The decay rate denotes the disappearance of the corresponding catalytic intermediate complexes.

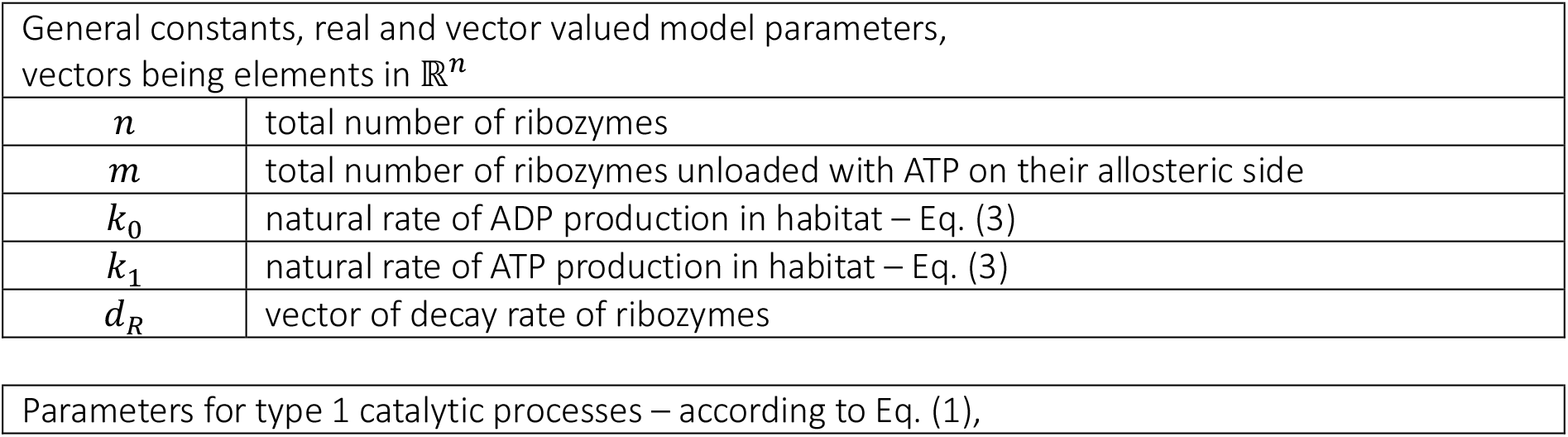
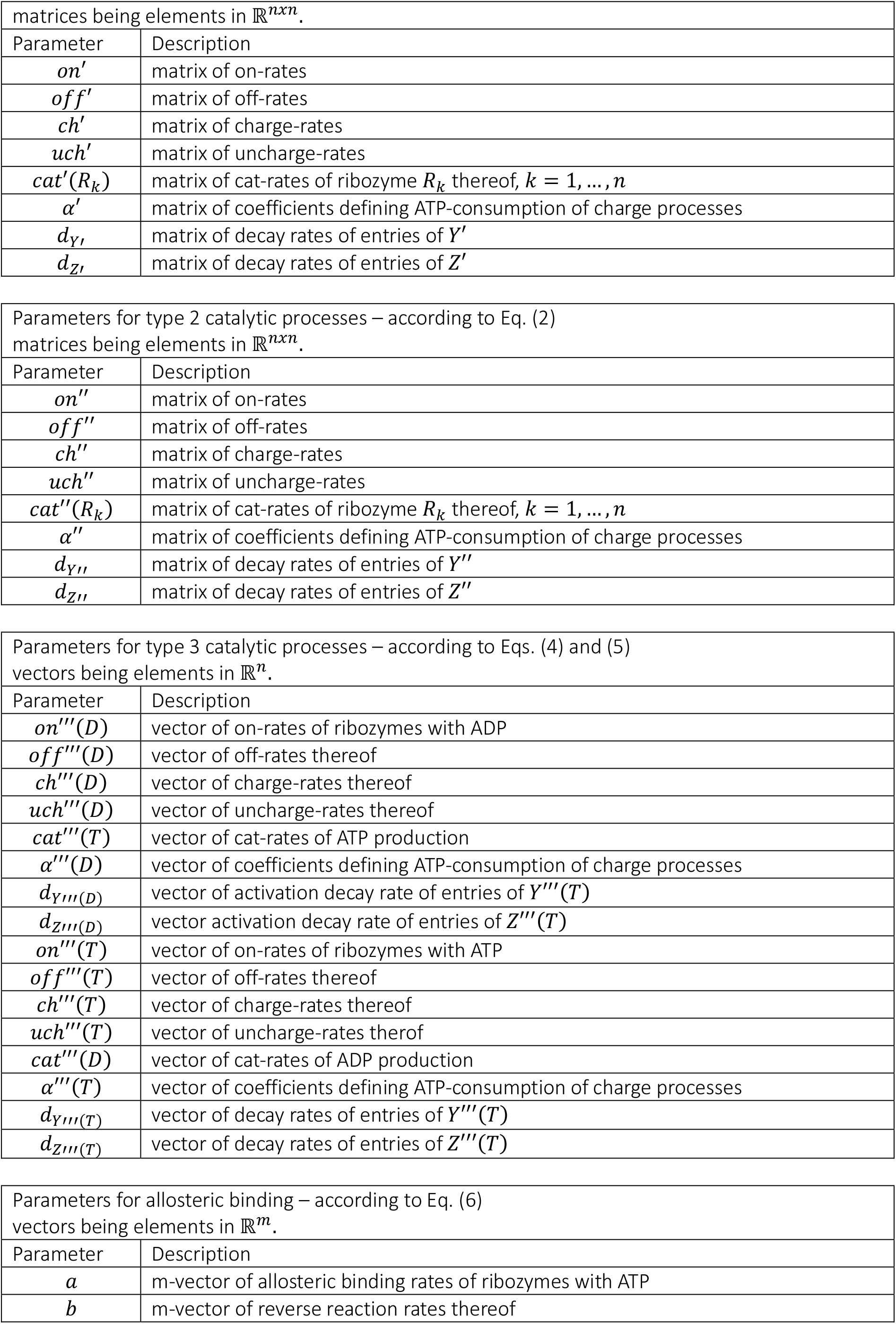

| General constants, real and vector valued model parameters,<br>vectors being elements in $\mathbb{R}^n$ | |
| --- | --- |
| $n$ | total number of ribozymes |
| $m$ | total number of ribozymes unloaded with ATP on their allosteric side |
| $k_0$ | natural rate of ADP production in habitat – Eq. (3) |
| $k_1$ | natural rate of ATP production in habitat – Eq. (3) |
| $d_R$ | vector of decay rate of ribozymes |

|  |
| --- |
| Parameters for type 1 catalytic processes – according to Eq. (1), |

| matrices being elements in $\mathbb{R}^{n \times n}$. | |
| --- | --- |
| Parameter | Description |
| $on'$ | matrix of on-rates |
| $off'$ | matrix of off-rates |
| $ch'$ | matrix of charge-rates |
| $uch'$ | matrix of uncharge-rates |
| $cat'(R_k)$ | matrix of cat-rates of ribozyme $R_k$ thereof, $k = 1, \dots, n$ |
| $\alpha'$ | matrix of coefficients defining ATP-consumption of charge processes |
| $d_{Y'}$ | matrix of decay rates of entries of $Y'$ |
| $d_{Z'}$ | matrix of decay rates of entries of $Z'$ |

| Parameters for type 2 catalytic processes – according to Eq. (2)<br>matrices being elements in $\mathbb{R}^{n \times n}$. | |
| --- | --- |
| Parameter | Description |
| $on''$ | matrix of on-rates |
| $off''$ | matrix of off-rates |
| $ch''$ | matrix of charge-rates |
| $uch''$ | matrix of uncharge-rates |
| $cat''(R_k)$ | matrix of cat-rates of ribozyme $R_k$ thereof, $k = 1, \dots, n$ |
| $\alpha''$ | matrix of coefficients defining ATP-consumption of charge processes |
| $d_{Y''}$ | matrix of decay rates of entries of $Y''$ |
| $d_{Z''}$ | matrix of decay rates of entries of $Z''$ |

| Parameters for type 3 catalytic processes – according to Eqs. (4) and (5)<br>vectors being elements in $\mathbb{R}^n$. | |
| --- | --- |
| Parameter | Description |
| $on'''(D)$ | vector of on-rates of ribozymes with ADP |
| $off'''(D)$ | vector of off-rates thereof |
| $ch'''(D)$ | vector of charge-rates thereof |
| $uch'''(D)$ | vector of uncharge-rates thereof |
| $cat'''(T)$ | vector of cat-rates of ATP production |
| $\alpha'''(D)$ | vector of coefficients defining ATP-consumption of charge processes |
| $d_{Y'''(D)}$ | vector of activation decay rate of entries of $Y'''(T)$ |
| $d_{Z'''(D)}$ | vector activation decay rate of entries of $Z'''(T)$ |
| $on'''(T)$ | vector of on-rates of ribozymes with ATP |
| $off'''(T)$ | vector of off-rates thereof |
| $ch'''(T)$ | vector of charge-rates thereof |
| $uch'''(T)$ | vector of uncharge-rates thereof |
| $cat'''(D)$ | vector of cat-rates of ADP production |
| $\alpha'''(T)$ | vector of coefficients defining ATP-consumption of charge processes |
| $d_{Y'''(T)}$ | vector of decay rates of entries of $Y'''(T)$ |
| $d_{Z'''(T)}$ | vector of decay rates of entries of $Z'''(T)$ |

| Parameters for allosteric binding – according to Eq. (6)<br>vectors being elements in $\mathbb{R}^m$. | |
| --- | --- |
| Parameter | Description |
| $a$ | m-vector of allosteric binding rates of ribozymes with ATP |
| $b$ | m-vector of reverse reaction rates thereof |

#### 1.2 Concrete parametrizations

##### 1.2.1 Model calculations for Fig. 1

Fig. 1 is a pure schematic exhibit of a possible evolution of the ATP-ADP cycle. Therefore parametrization is irrelevant here.

##### 1.2.2 Model calculations for Figs 2

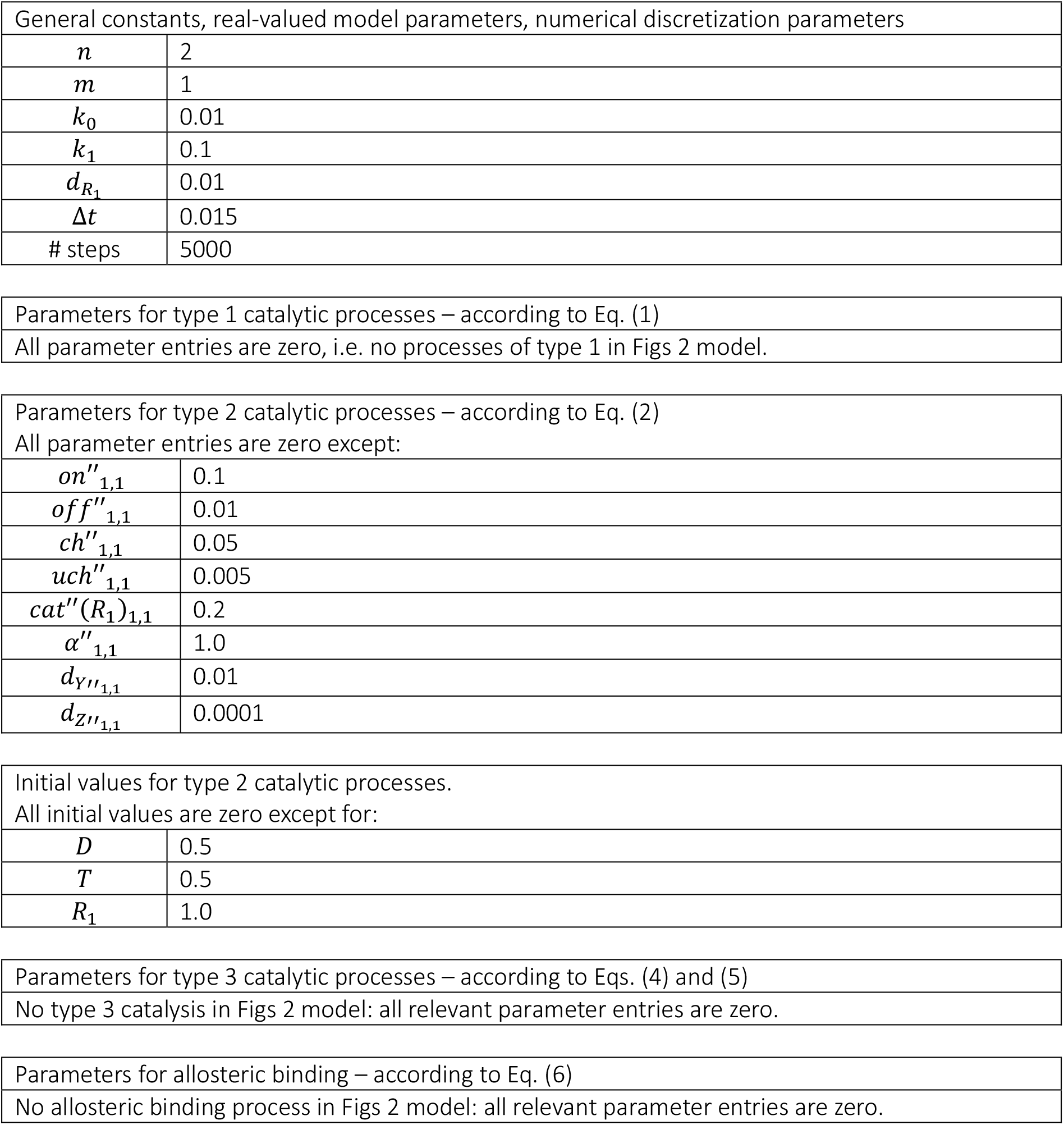

| General constants, real-valued model parameters, numerical discretization parameters |  |
| --- | --- |
| $n$ | 2 |
| $m$ | 1 |
| $k_0$ | 0.01 |
| $k_1$ | 0.1 |
| $d_{R_1}$ | 0.01 |
| $\Delta t$ | 0.015 |
| # steps | 5000 |

|  |
| --- |
| Parameters for type 1 catalytic processes – according to Eq. (1) |
| All parameter entries are zero, i.e. no processes of type 1 in Figs 2 model. |
| Parameters for type 2 catalytic processes – according to Eq. (2) |  |
| All parameter entries are zero except: |  |
| $on''_{1,1}$ | 0.1 |
| $off''_{1,1}$ | 0.01 |
| $ch''_{1,1}$ | 0.05 |
| $uch''_{1,1}$ | 0.005 |
| $cat''(R_1)_{1,1}$ | 0.2 |
| $\alpha''_{1,1}$ | 1.0 |
| $d_{Y''_{1,1}}$ | 0.01 |
| $d_{Z''_{1,1}}$ | 0.0001 |
| Initial values for type 2 catalytic processes.<br>All initial values are zero except for: |  |
| $D$ | 0.5 |
| $T$ | 0.5 |
| $R_1$ | 1.0 |

|  |
| --- |
| Parameters for type 3 catalytic processes – according to Eqs. (4) and (5) |
| No type 3 catalysis in Figs 2 model: all relevant parameter entries are zero. |
| Parameters for allosteric binding – according to Eq. (6) |
| No allosteric binding process in Figs 2 model: all relevant parameter entries are zero. |

##### 1.2.3 Model calculations for Figs 3

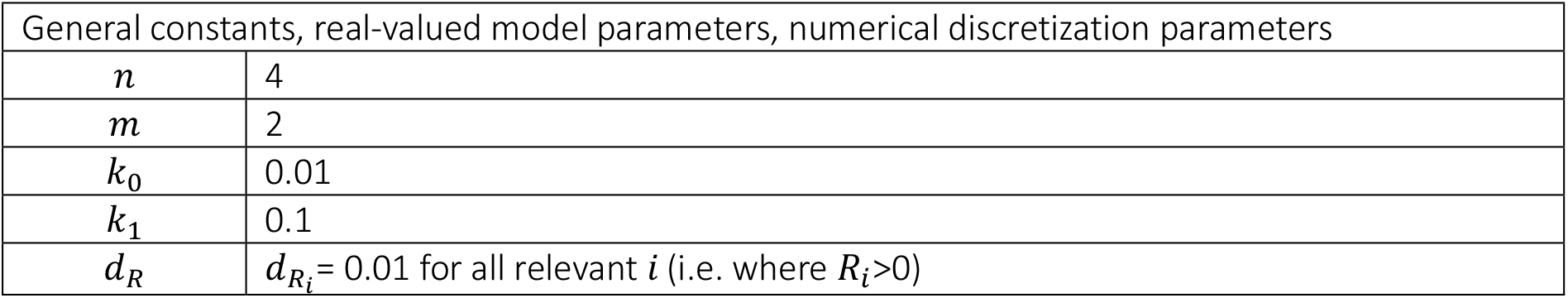
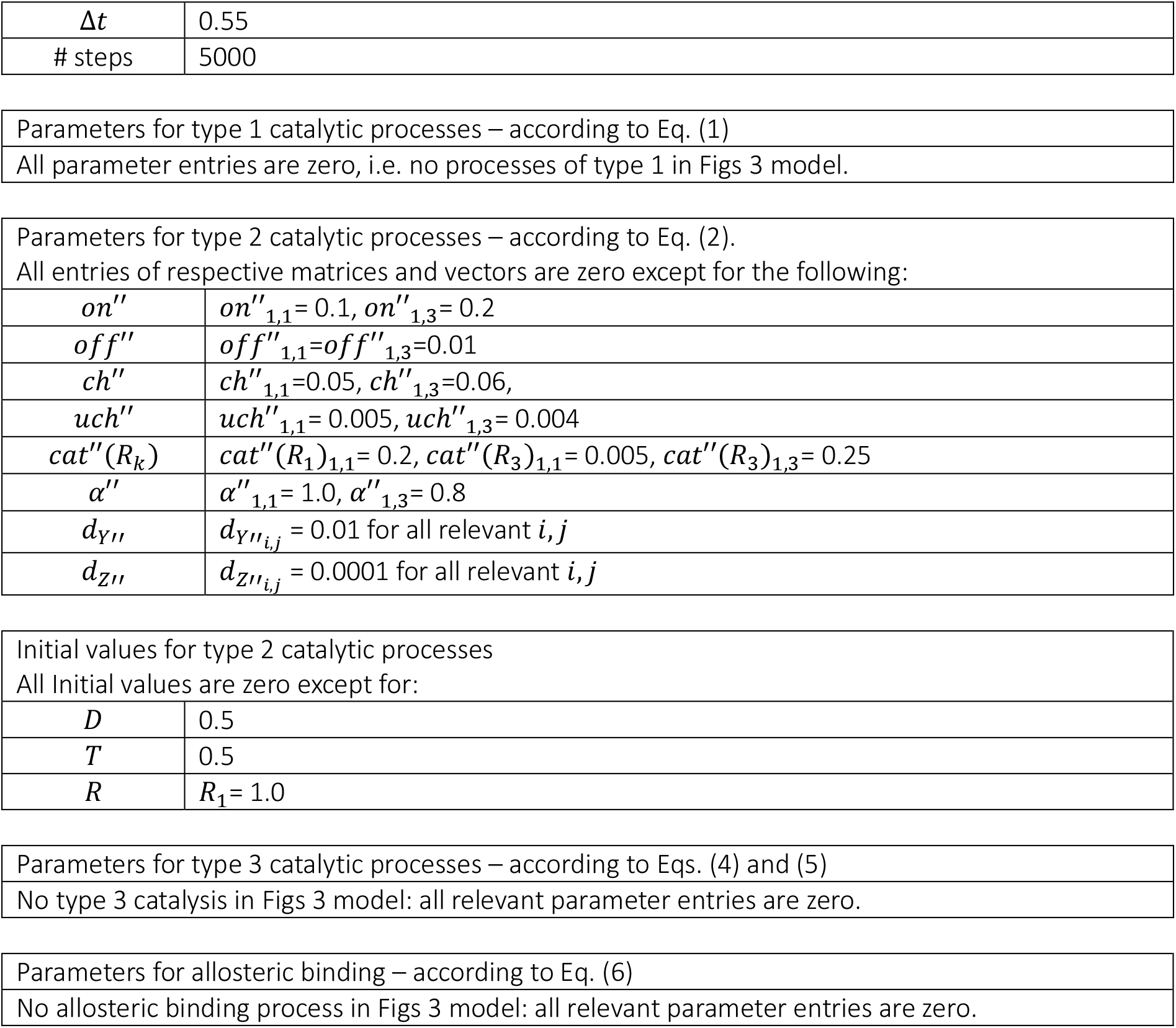

| General constants, real-valued model parameters, numerical discretization parameters |  |
| --- | --- |
| $n$ | 4 |
| $m$ | 2 |
| $k_0$ | 0.01 |
| $k_1$ | 0.1 |
| $d_R$ | $d_{R_i} = 0.01$ for all relevant $i$ (i.e. where $R_i > 0$) |
| $\Delta t$ | 0.55 |
| # steps | 5000 |
| $on''$ | $on''_{1,1}=0.1, on''_{1,3}=0.2$ |
| $off''$ | $off''_{1,1}=off''_{1,3}=0.01$ |
| $ch''$ | $ch''_{1,1}=0.05, ch''_{1,3}=0.06,$ |
| $uch''$ | $uch''_{1,1}=0.005, uch''_{1,3}=0.004$ |
| $cat''(R_k)$ | $cat''(R_1)_{1,1}=0.2, cat''(R_3)_{1,1}=0.005, cat''(R_3)_{1,3}=0.25$ |
| $\alpha''$ | $\alpha''_{1,1}=1.0, \alpha''_{1,3}=0.8$ |
| $d_{Y''}$ | $d_{Y''_{i,j}}=0.01$ for all relevant $i, j$ |
| $d_{Z''}$ | $d_{Z''_{i,j}}=0.0001$ for all relevant $i, j$ |
| $D$ | 0.5 |
| $T$ | 0.5 |
| $R$ | $R_1=1.0$ |

##### 1.2.4 Model calculations for Figs 4

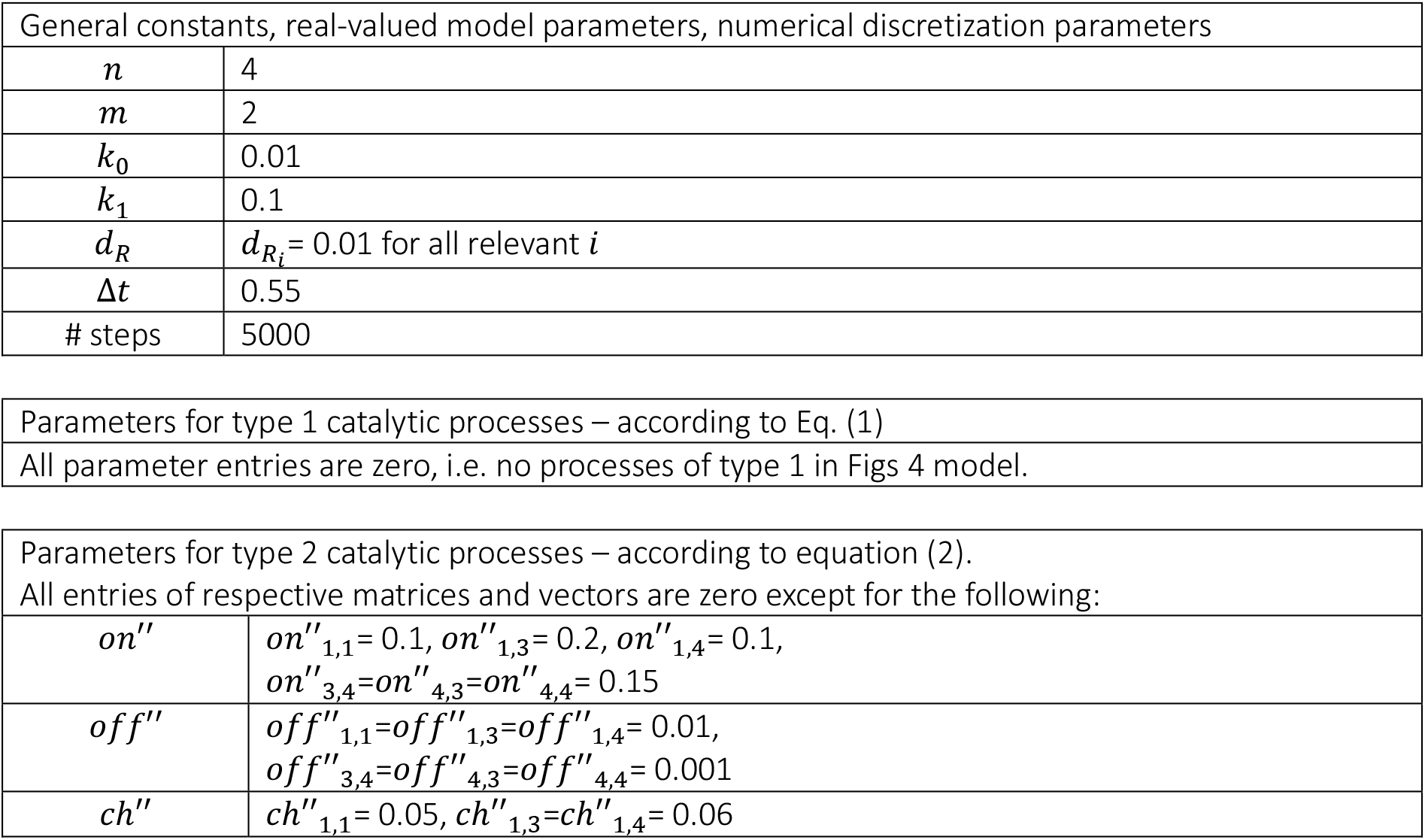
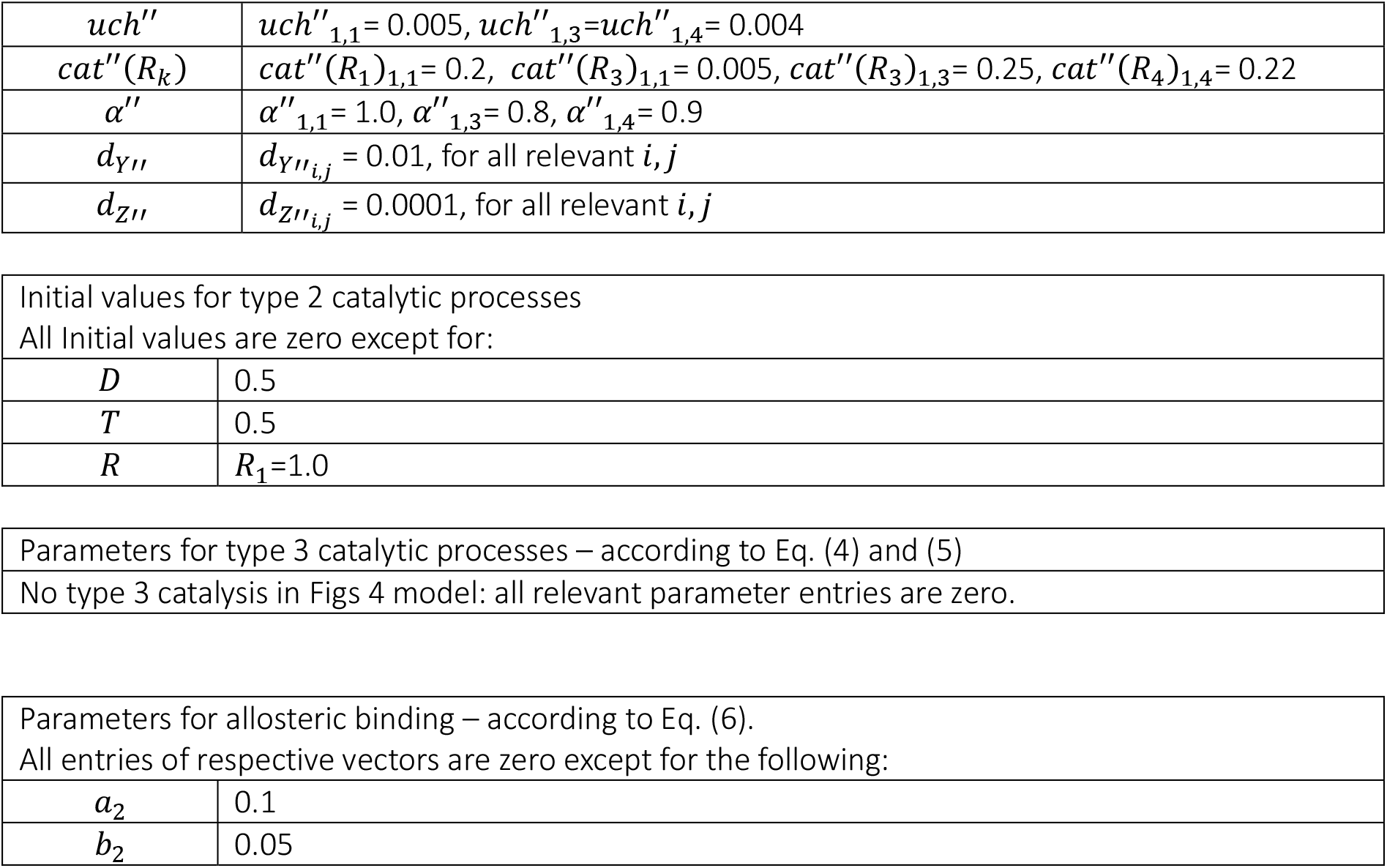

|  |  |
| --- | --- |
| $n$ | 4 |
| $m$ | 2 |
| $k_0$ | 0.01 |
| $k_1$ | 0.1 |
| $d_R$ | $d_{R_i}=0.01$ for all relevant $i$ |
| $\Delta t$ | 0.55 |
| # steps | 5000 |
| $on''$ | $on''_{1,1}=0.1, on''_{1,3}=0.2, on''_{1,4}=0.1,$<br>$on''_{3,4}=on''_{4,3}=on''_{4,4}=0.15$ |
| $off''$ | $off''_{1,1}=off''_{1,3}=off''_{1,4}=0.01,$<br>$off''_{3,4}=off''_{4,3}=off''_{4,4}=0.001$ |
| $ch''$ | $ch''_{1,1}=0.05, ch''_{1,3}=ch''_{1,4}=0.06$ |
| $uch''$ | $uch''_{1,1} = 0.005, uch''_{1,3}=uch''_{1,4} = 0.004$ |
| $cat''(R_k)$ | $cat''(R_1)_{1,1} = 0.2, cat''(R_3)_{1,1} = 0.005, cat''(R_3)_{1,3} = 0.25, cat''(R_4)_{1,4} = 0.22$ |
| $\alpha''$ | $\alpha''_{1,1} = 1.0, \alpha''_{1,3} = 0.8, \alpha''_{1,4} = 0.9$ |
| $d_{Y''}$ | $d_{Y''_{i,j}} = 0.01$, for all relevant $i, j$ |
| $d_{Z''}$ | $d_{Z''_{i,j}} = 0.0001$, for all relevant $i, j$ |
| $a_2$ | 0.1 |
| $b_2$ | 0.05 |

##### 1.2.5 Model calculations for Figs 5

Figs 5 use the same parametrization as Figs 4, except the following parameters:

|  |  |
| --- | --- |
| $k_0$ | 0.02, double rate as compared to Figs 4 |
| $k_1$ | 0.2, double rate as compared to Figs 4 |

##### 1.2.6 Model calculations for Figs 6

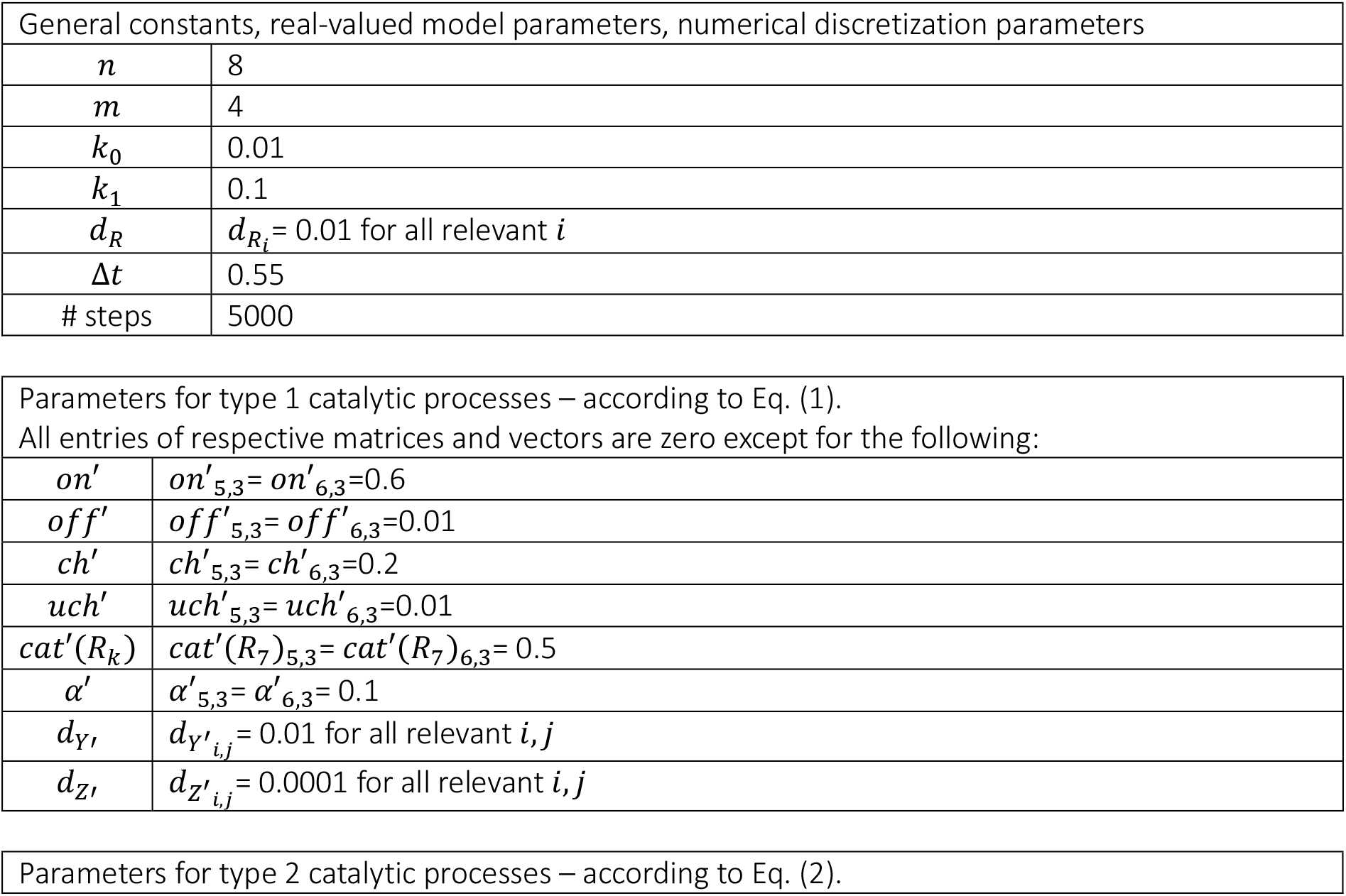
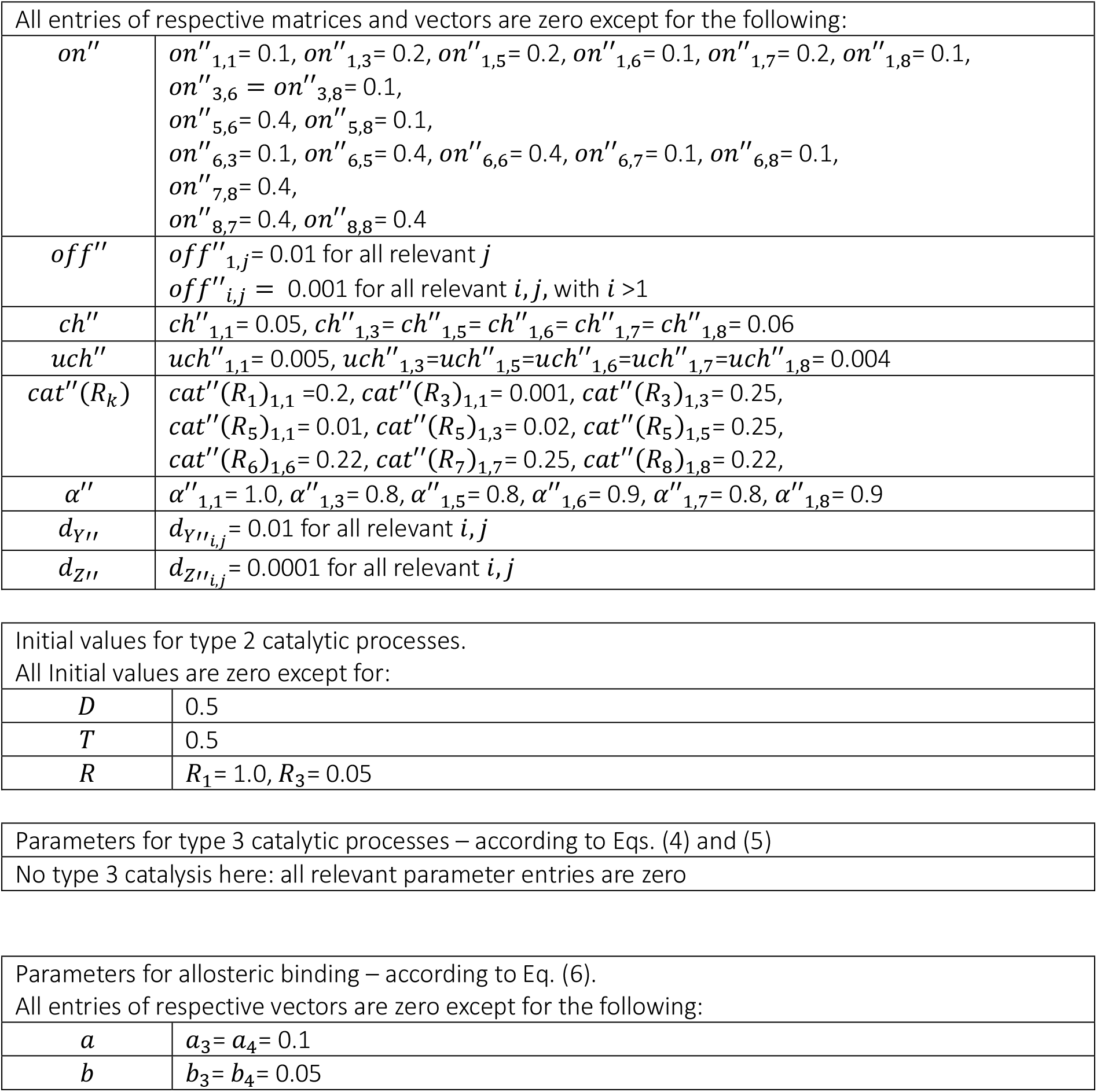

|  |  |
| --- | --- |
| $n$ | 8 |
| $m$ | 4 |
| $k_0$ | 0.01 |
| $k_1$ | 0.1 |
| $d_R$ | $d_{R_i} = 0.01$ for all relevant $i$ |
| $\Delta t$ | 0.55 |
| # steps | 5000 |
| $on'$ | $on'_{5,3} = on'_{6,3} = 0.6$ |
| $off'$ | $off'_{5,3} = off'_{6,3} = 0.01$ |
| $ch'$ | $ch'_{5,3} = ch'_{6,3} = 0.2$ |
| $uch'$ | $uch'_{5,3} = uch'_{6,3} = 0.01$ |
| $cat'(R_k)$ | $cat'(R_7)_{5,3} = cat'(R_7)_{6,3} = 0.5$ |
| $\alpha'$ | $\alpha'_{5,3} = \alpha'_{6,3} = 0.1$ |
| $d_{Y'}$ | $d_{Y'_{i,j}} = 0.01$ for all relevant $i, j$ |
| $d_{Z'}$ | $d_{Z'_{i,j}} = 0.0001$ for all relevant $i, j$ |
| All entries of respective matrices and vectors are zero except for the following: |  |
| $on''$ | $on''_{1,1}=0.1, on''_{1,3}=0.2, on''_{1,5}=0.2, on''_{1,6}=0.1, on''_{1,7}=0.2, on''_{1,8}=0.1,$<br>$on''_{3,6}=on''_{3,8}=0.1,$<br>$on''_{5,6}=0.4, on''_{5,8}=0.1,$<br>$on''_{6,3}=0.1, on''_{6,5}=0.4, on''_{6,6}=0.4, on''_{6,7}=0.1, on''_{6,8}=0.1,$<br>$on''_{7,8}=0.4,$<br>$on''_{8,7}=0.4, on''_{8,8}=0.4$ |
| $off''$ | $off''_{1,j}=0.01$ for all relevant $j$<br>$off''_{i,j}=0.001$ for all relevant $i,j$, with $i>1$ |
| $ch''$ | $ch''_{1,1}=0.05, ch''_{1,3}=ch''_{1,5}=ch''_{1,6}=ch''_{1,7}=ch''_{1,8}=0.06$ |
| $uch''$ | $uch''_{1,1}=0.005, uch''_{1,3}=uch''_{1,5}=uch''_{1,6}=uch''_{1,7}=uch''_{1,8}=0.004$ |
| $cat''(R_k)$ | $cat''(R_1)_{1,1}=0.2, cat''(R_3)_{1,1}=0.001, cat''(R_3)_{1,3}=0.25,$<br>$cat''(R_5)_{1,1}=0.01, cat''(R_5)_{1,3}=0.02, cat''(R_5)_{1,5}=0.25,$<br>$cat''(R_6)_{1,6}=0.22, cat''(R_7)_{1,7}=0.25, cat''(R_8)_{1,8}=0.22,$ |
| $\alpha''$ | $\alpha''_{1,1}=1.0, \alpha''_{1,3}=0.8, \alpha''_{1,5}=0.8, \alpha''_{1,6}=0.9, \alpha''_{1,7}=0.8, \alpha''_{1,8}=0.9$ |
| $d_{Y''}$ | $d_{Y''_{i,j}}=0.01$ for all relevant $i,j$ |
| $d_{Z''}$ | $d_{Z''_{i,j}}=0.0001$ for all relevant $i,j$ |
| Initial values for type 2 catalytic processes.<br>All Initial values are zero except for: |  |
| $D$ | 0.5 |
| $T$ | 0.5 |
| $R$ | $R_1=1.0, R_3=0.05$ |
| Parameters for type 3 catalytic processes – according to Eqs. (4) and (5) |  |
| No type 3 catalysis here: all relevant parameter entries are zero |  |
| Parameters for allosteric binding – according to Eq. (6).<br>All entries of respective vectors are zero except for the following: |  |
| $a$ | $a_3=a_4=0.1$ |
| $b$ | $b_3=b_4=0.05$ |

##### 1.2.7 Model calculations for Figs 7

Figs 7 follow the same parametrization as Figs 6, except for the following parameters:

|  |  |
| --- | --- |
| $k_0$ | 0.03, triple rate as compared to Figs 4 |
| $k_1$ | 0.3, triple rate as compares to Figs 4 |

##### 1.2.8 Model calculations for Figs 8

|  |  |
| --- | --- |
| General constants, real-valued model parameters, numerical discretization parameters |  |
| $n$ | 4 |
| $m$ | 2 |
| $k_0$ | 0.01 |
| $k_1$ | 0.1 |
| $d_R$ | $d_{R_i}=0.01$ for all relevant $i$ |
| $\Delta t$ | 0.55 |
| # steps | 5000 |
| Parameters for type 1 catalytic processes – according to Eq. (1) |  |
| All parameter entries are zero, i.e. no processes of type 1 in Figs 8 model. |  |
| Parameters for type 2 catalytic processes – according to Eq. (2). |  |
| All entries of respective matrices and vectors are zero except for the following: |  |
| $on''$ | $on''_{1,1} = 0.1, on''_{1,3} = 0.2, on''_{1,4} = 0.1,$<br>$on''_{3,4} = 0.15,$<br>$on''_{4,3} = on''_{4,4} = 0.15$ |
| $off''$ | $off''_{1,j} = 0.01$ for all relevant $j$,<br>$off''_{i,j} = 0.001$ for all relevant $i, j$, with $i > 1$ |
| $ch''$ | $ch''_{1,1} = 0.05, ch''_{1,3} = ch''_{1,4} = 0.06$ |
| $uch''$ | $uch''_{1,1} = 0.005, uch''_{1,3} = uch''_{1,4} = 0.004$ |
| $cat''(R_k)$ | $cat''(R_1)_{1,1} = 0.2,$<br>$cat''(R_3)_{1,1} = 0.001, cat''(R_3)_{1,3} = 0.25,$<br>$cat''(R_4)_{1,4} = 0.22,$ |
| $\alpha''$ | $\alpha''_{1,1} = 1.0, \alpha''_{1,3} = 0.8, \alpha''_{1,4} = 0.9$ |
| $d_{Y''}$ | $d_{Y''_{i,j}} = 0.01$ for all relevant $i, j$ |
| $d_{Z''}$ | $d_{Z''_{i,j}} = 0.0001$ for all relevant $i, j$ |
| Initial values for type 2 catalytic processes. |  |
| All Initial values are zero except for: |  |
| $D$ | 0.5 |
| $T$ | 0.5 |
| $R$ | $R_1 = 1.0, R_3 = 0.05$ |

| Parameters for type 3 catalytic processes – according to Eqs. (4) and (5) |  |
| --- | --- |
| All entries of respective vectors below are zero except the following: |  |
| Parameter | Description |
| $on'''(D)_3$ | 0.2 |
| $off'''(D)_3$ | 0.01 |
| $ch'''(D)_3$ | 0.2 |
| $uch'''(D)_3$ | 0.01 |
| $cat'''(T)_3$ | 1.0 |
| $\alpha'''(D)_3$ | 0.1 |
| $d_{Y'''(D)_3}$ | 0.01 |
| $d_{Z'''(D)_3}$ | 0.01 |
| $on'''(T)_3$ | 0.2 |
| $off'''(T)_3$ | 0.01 |
| $ch'''(T)_3$ | 0.2 |
| $uch'''(T)_3$ | 0.01 |
| $cat'''(D)_3$ | 1.0 |
| $\alpha'''(T)_3$ | 0.1 |
| $d_{Y'''(T)_3}$ | 0.01 |
| $d_{Z'''(T)_3}$ | 0.01 |
| Parameters for allosteric binding – according to Eq. (6) |  |
| No allosteric binding process in Figs 8 model: all relevant parameter entries are zero. |  |

##### 1.2.9 Model calculations for Figs 9

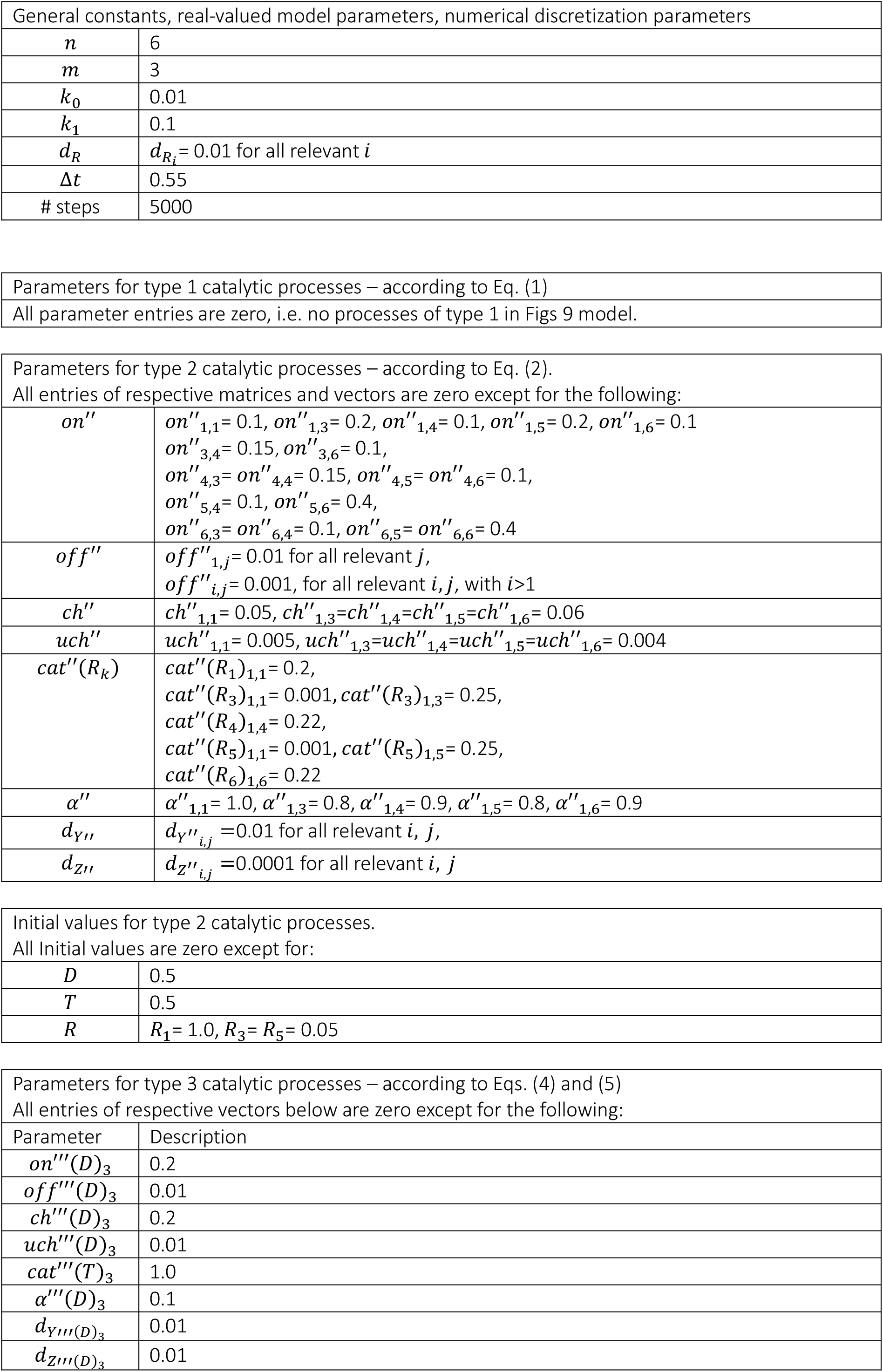
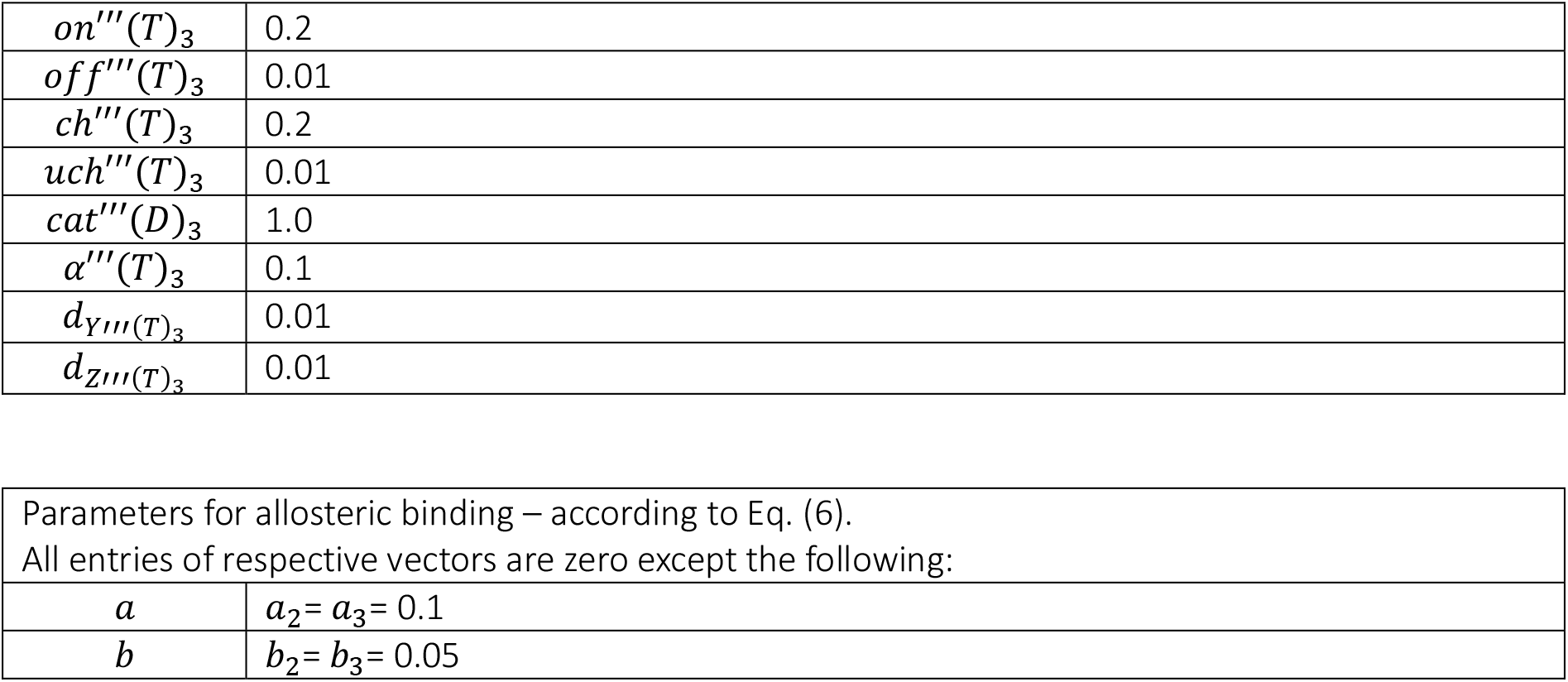

| General constants, real-valued model parameters, numerical discretization parameters |  |
| --- | --- |
| $n$ | 6 |
| $m$ | 3 |
| $k_0$ | 0.01 |
| $k_1$ | 0.1 |
| $d_R$ | $d_{R_i} = 0.01$ for all relevant $i$ |
| $\Delta t$ | 0.55 |
| # steps | 5000 |

|  |
| --- |
| Parameters for type 1 catalytic processes – according to Eq. (1) |
| All parameter entries are zero, i.e. no processes of type 1 in Figs 9 model. |
| Parameters for type 2 catalytic processes – according to Eq. (2).<br>All entries of respective matrices and vectors are zero except for the following: |  |
| $on''$ | $on''_{1,1} = 0.1, on''_{1,3} = 0.2, on''_{1,4} = 0.1, on''_{1,5} = 0.2, on''_{1,6} = 0.1$<br>$on''_{3,4} = 0.15, on''_{3,6} = 0.1,$<br>$on''_{4,3} = on''_{4,4} = 0.15, on''_{4,5} = on''_{4,6} = 0.1,$<br>$on''_{5,4} = 0.1, on''_{5,6} = 0.4,$<br>$on''_{6,3} = on''_{6,4} = 0.1, on''_{6,5} = on''_{6,6} = 0.4$ |
| $off''$ | $off''_{1,j} = 0.01$ for all relevant $j$,<br>$off''_{i,j} = 0.001$, for all relevant $i, j$, with $i > 1$ |
| $ch''$ | $ch''_{1,1} = 0.05, ch''_{1,3} = ch''_{1,4} = ch''_{1,5} = ch''_{1,6} = 0.06$ |
| $uch''$ | $uch''_{1,1} = 0.005, uch''_{1,3} = uch''_{1,4} = uch''_{1,5} = uch''_{1,6} = 0.004$ |
| $cat''(R_k)$ | $cat''(R_1)_{1,1} = 0.2,$<br>$cat''(R_3)_{1,1} = 0.001, cat''(R_3)_{1,3} = 0.25,$<br>$cat''(R_4)_{1,4} = 0.22,$<br>$cat''(R_5)_{1,1} = 0.001, cat''(R_5)_{1,5} = 0.25,$<br>$cat''(R_6)_{1,6} = 0.22$ |
| $\alpha''$ | $\alpha''_{1,1} = 1.0, \alpha''_{1,3} = 0.8, \alpha''_{1,4} = 0.9, \alpha''_{1,5} = 0.8, \alpha''_{1,6} = 0.9$ |
| $d_{Y''}$ | $d_{Y''_{i,j}} = 0.01$ for all relevant $i, j$, |
| $d_{Z''}$ | $d_{Z''_{i,j}} = 0.0001$ for all relevant $i, j$ |
| Initial values for type 2 catalytic processes.<br>All Initial values are zero except for: |  |
| $D$ | 0.5 |
| $T$ | 0.5 |
| $R$ | $R_1 = 1.0, R_3 = R_5 = 0.05$ |

| Parameters for type 3 catalytic processes – according to Eqs. (4) and (5)<br>All entries of respective vectors below are zero except for the following: |  |
| --- | --- |
| Parameter | Description |
| $on'''(D)_3$ | 0.2 |
| $off'''(D)_3$ | 0.01 |
| $ch'''(D)_3$ | 0.2 |
| $uch'''(D)_3$ | 0.01 |
| $cat'''(T)_3$ | 1.0 |
| $\alpha'''(D)_3$ | 0.1 |
| $d_{Y'''(D)_3}$ | 0.01 |
| $d_{Z'''(D)_3}$ | 0.01 |
| $on'''(T)_3$ | 0.2 |
| $off'''(T)_3$ | 0.01 |
| $ch'''(T)_3$ | 0.2 |
| $uch'''(T)_3$ | 0.01 |
| $cat'''(D)_3$ | 1.0 |
| $\alpha'''(T)_3$ | 0.1 |
| $d_{Y'''(T)_3}$ | 0.01 |
| $d_{Z'''(T)_3}$ | 0.01 |
| $a$ | $a_2 = a_3 = 0.1$ |
| $b$ | $b_2 = b_3 = 0.05$ |

##### 1.2.10 Model calculations for Figs 10

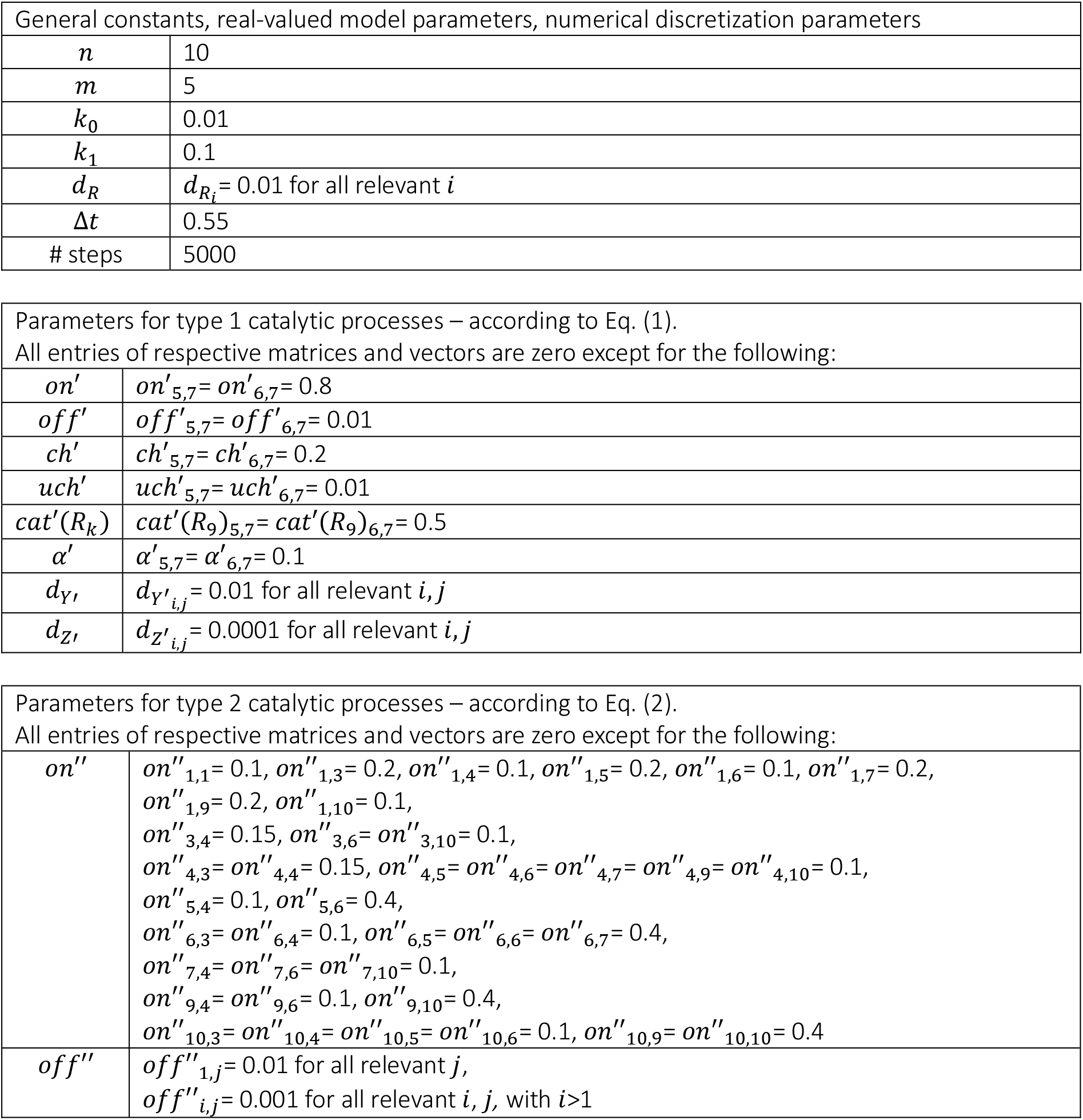
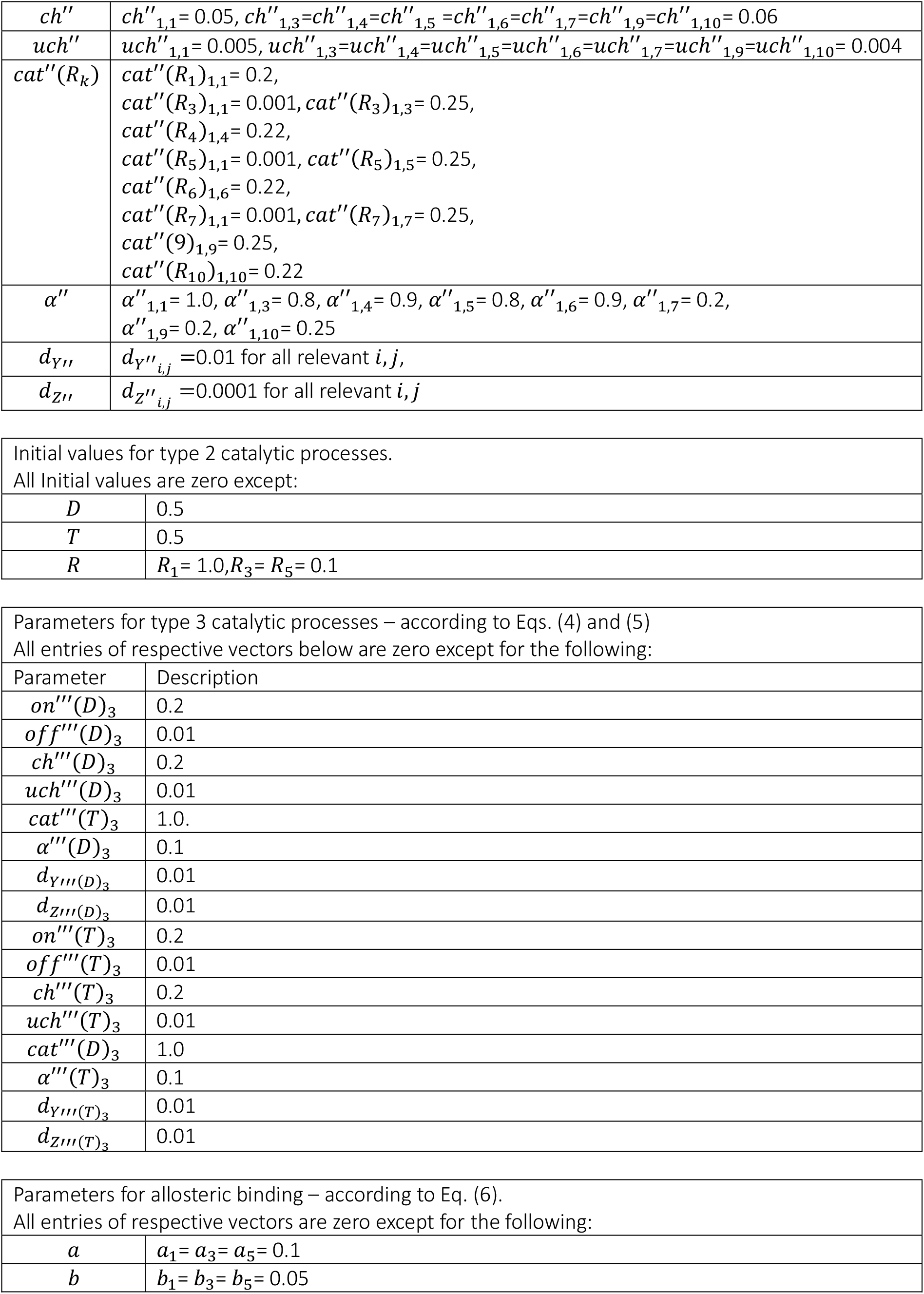

| General constants, real-valued model parameters, numerical discretization parameters |  |
| --- | --- |
| $n$ | 10 |
| $m$ | 5 |
| $k_0$ | 0.01 |
| $k_1$ | 0.1 |
| $d_R$ | $d_{R_i} = 0.01$ for all relevant $i$ |
| $\Delta t$ | 0.55 |
| # steps | 5000 |
| $on'$ | $on'_{5,7} = on'_{6,7} = 0.8$ |
| $off'$ | $off'_{5,7} = off'_{6,7} = 0.01$ |
| $ch'$ | $ch'_{5,7} = ch'_{6,7} = 0.2$ |
| $uch'$ | $uch'_{5,7} = uch'_{6,7} = 0.01$ |
| $cat'(R_k)$ | $cat'(R_9)_{5,7} = cat'(R_9)_{6,7} = 0.5$ |
| $\alpha'$ | $\alpha'_{5,7} = \alpha'_{6,7} = 0.1$ |
| $d_{Y'}$ | $d_{Y'_{i,j}} = 0.01$ for all relevant $i, j$ |
| $d_{Z'}$ | $d_{Z'_{i,j}} = 0.0001$ for all relevant $i, j$ |
| $on''$ | $on''_{1,1} = 0.1, on''_{1,3} = 0.2, on''_{1,4} = 0.1, on''_{1,5} = 0.2, on''_{1,6} = 0.1, on''_{1,7} = 0.2,$<br>$on''_{1,9} = 0.2, on''_{1,10} = 0.1,$<br>$on''_{3,4} = 0.15, on''_{3,6} = on''_{3,10} = 0.1,$<br>$on''_{4,3} = on''_{4,4} = 0.15, on''_{4,5} = on''_{4,6} = on''_{4,7} = on''_{4,9} = on''_{4,10} = 0.1,$<br>$on''_{5,4} = 0.1, on''_{5,6} = 0.4,$<br>$on''_{6,3} = on''_{6,4} = 0.1, on''_{6,5} = on''_{6,6} = on''_{6,7} = 0.4,$<br>$on''_{7,4} = on''_{7,6} = on''_{7,10} = 0.1,$<br>$on''_{9,4} = on''_{9,6} = 0.1, on''_{9,10} = 0.4,$<br>$on''_{10,3} = on''_{10,4} = on''_{10,5} = on''_{10,6} = 0.1, on''_{10,9} = on''_{10,10} = 0.4$ |
| $off''$ | $off''_{1,j} = 0.01$ for all relevant $j,$<br>$off''_{i,j} = 0.001$ for all relevant $i, j,$ with $i > 1$ |
| $ch''$ | $ch''_{1,1} = 0.05, ch''_{1,3}=ch''_{1,4}=ch''_{1,5}=ch''_{1,6}=ch''_{1,7}=ch''_{1,9}=ch''_{1,10} = 0.06$ |
| $uch''$ | $uch''_{1,1} = 0.005, uch''_{1,3}=uch''_{1,4}=uch''_{1,5}=uch''_{1,6}=uch''_{1,7}=uch''_{1,9}=uch''_{1,10} = 0.004$ |
| $cat''(R_k)$ | $cat''(R_1)_{1,1} = 0.2,$<br>$cat''(R_3)_{1,1} = 0.001, cat''(R_3)_{1,3} = 0.25,$<br>$cat''(R_4)_{1,4} = 0.22,$<br>$cat''(R_5)_{1,1} = 0.001, cat''(R_5)_{1,5} = 0.25,$<br>$cat''(R_6)_{1,6} = 0.22,$<br>$cat''(R_7)_{1,1} = 0.001, cat''(R_7)_{1,7} = 0.25,$<br>$cat''(9)_{1,9} = 0.25,$<br>$cat''(R_{10})_{1,10} = 0.22$ |
| $\alpha''$ | $\alpha''_{1,1} = 1.0, \alpha''_{1,3} = 0.8, \alpha''_{1,4} = 0.9, \alpha''_{1,5} = 0.8, \alpha''_{1,6} = 0.9, \alpha''_{1,7} = 0.2,$<br>$\alpha''_{1,9} = 0.2, \alpha''_{1,10} = 0.25$ |
| $d_{Y''}$ | $d_{Y''_{i,j}} = 0.01$ for all relevant $i, j,$ |
| $d_{Z''}$ | $d_{Z''_{i,j}} = 0.0001$ for all relevant $i, j$ |
| $D$ | 0.5 |
| $T$ | 0.5 |
| $R$ | $R_1 = 1.0, R_3 = R_5 = 0.1$ |

Parameters for type 3 catalytic processes – according to Eqs. (4) and (5) All entries of respective vectors below are zero except for the following:
| Parameter | Description |
| --- | --- |
| $on'''(D)_3$ | 0.2 |
| $off'''(D)_3$ | 0.01 |
| $ch'''(D)_3$ | 0.2 |
| $uch'''(D)_3$ | 0.01 |
| $cat'''(T)_3$ | 1.0. |
| $\alpha'''(D)_3$ | 0.1 |
| $d_{Y'''}(D)_3$ | 0.01 |
| $d_{Z'''}(D)_3$ | 0.01 |
| $on'''(T)_3$ | 0.2 |
| $off'''(T)_3$ | 0.01 |
| $ch'''(T)_3$ | 0.2 |
| $uch'''(T)_3$ | 0.01 |
| $cat'''(D)_3$ | 1.0 |
| $\alpha'''(T)_3$ | 0.1 |
| $d_{Y'''}(T)_3$ | 0.01 |
| $d_{Z'''}(T)_3$ | 0.01 |
| $a$ | $a_1 = a_3 = a_5 = 0.1$ |
| $b$ | $b_1 = b_3 = b_5 = 0.05$ |

## References

Anet, F.A.L. The place of metabolism in the origin of life. Curr. Opin. Chem. Biol. 2004, 8, 654–659. 10.1016/j.cbpa.2004.10.005.

Briones, C.; Stich, M.; Manrubia, S.C. The dawn of the RNA world: toward functional complexity through ligation of random RNA oligomers. RNA 2009, 15, 743–749. 10.1261/rna.1488609.

Buss, L.W. The Evolution of Individuality. Princeton (NJ): Princeton University Press; 1987.

Conrad, B.; Iseli, C.; Pirovino, M. Energy-harnessing problem solving of primordial life. PLoS One 2023, 18, e0281661. 10.1371/journal.pone.0281661.

Fenton, A.W. Allostery: an illustrated definition for the ‘second secret of life’. Trends Biochem. Sci. 2008, 33, 420–425. 10.1016/j.tibs.2008.05.009.

Gilbert, W. Origin of life: the RNAworld. Nature 1986, 319, 618. 10.1038/319618a0

Gianni, E.; Kwok, S.L.Y.; Wan, C.J.K.; et al. A small polymerase ribozyme that can synthesize itself and its complementary strand. Science 2026, 391, 1022–1028. 10.1126/science.adt2760.

Hardie D.G., Scott, J.W., Pan, D.A., Hudson, E.R. Management of cellular energy by the AMP-activated protein kinase system. FEBS Lett. 2003, 546, 113–120. 10.1016/S0014-5793(03)00560-X.

Hayden, E.J.; Lehman, N. Self-assembly of a group I intron from inactive oligonucleotide fragments. Chem. Biol. 2006, 13, 909–918. 10.1016/j.chembiol.2006.06.014.

Iranzo, J.; Puigbò, P.; Lobkovsky, A.E.; Wolf, Y.I.; Koonin, E.V. Inevitability of genetic parasites. Genome Biol. Evol. 2016, 8, 2856–2869. 10.1093/gbe/evw193.

Kauffman, S.A. Autocatalytic sets of proteins. J. Theor. Biol. 1986, 119, 1–24. 10.1016/S0022-5193(86)80047-9.

Kirschning, A. On the evolutionary history of the twenty encoded amino acids. Chemistry 2022, 28, e202201419. 10.1002/chem.202201419.

Koonin, E.V.; Wolf, Y.I.; Katsnelson, M.I. Inevitability of the emergence and persistence of genetic parasites caused by evolutionary instability of parasite-free states. Biol. Direct 2017, 12, 31. 10.1186/s13062-017-0202-5.

Krakauer, D.C., Bertschinger, N., Olbrich, E., Ay, N., Flack, J.C. The information theory of individuality. Theory Biosci. 2020, 139, 209–223.

Macdonald, J.E.; Ashby, P.D. The molecular mechanism of ATP synthase constrains the evolutionary landscape of chemiosmosis. Biophys. J. 2025, 124, 2103–2119. 10.1016/j.bpj.2025.05.017.

Maynard Smith J., Szathmáry, E. The Major Transitions in Evolution. Oxford: W.H. Freeman (Spektrum); 1995.

Monod, J. Chance and Necessity: Essay on the Natural Philosophy of Modern Biology. Penguin Books, London, 1977.

Montserrat-Canals, M.; Cordara, G.; Krengel, U. Allostery. Q. Rev. Biophys. 2025, 58, e5. 10.1017/S0033583524000209.

Morowitz, H.J. A theory of biochemical organization, metabolic pathways, and evolution. Complexity 1999, 4, 39–53. 10.1002/(SICI)1099-0526(199907/08)4:6<39::AID-CPLX8>3.0.CO;2-2.

Nghe, P. A stepwise emergence of evolution in the RNA world. FEBS Lett. 2025, 599, 2706–2717. 10.1002/1873-3468.70065.

Nowak, M.A. Five rules for the evolution of cooperation. Science. 2006, 314, 1560–1563.

Papastavrou, N.; Horning, D.P.; Joyce, G.F. RNA-catalyzed evolution of catalytic RNA. Proc. Natl. Acad. Sci. U.S.A. 2024, 121, e2321592121. 10.1073/pnas.2321592121.

Pinna, S.; Kunz, C.; Halpern, A.; et al. A prebiotic basis for ATP as the universal energy currency. PLoS Biol. 2022, 20, e3001437. 10.1371/journal.pbio.3001437.

Pirovino, M.; Iseli, C.; Curran, J.A.; Conrad, B. Biomathematical enzyme kinetics model of prebiotic autocatalytic RNA networks. PLoS Comput. Biol. 2025, 21, e1012162. 10.1371/journal.pcbi.1012162.

Pross, A. Causation and the origin of life. Metabolism or replication first? Orig. Life Evol. Biosph. 2004, 34, 307–321. 10.1023/B:ORIG.0000016446.51012.bc.

Shah V., de Bouter J., Pauli Q., Tupper A.S., Higgs P.G. Survival of RNA replicators is much easier in protocells than in surface-based, spatial systems. Life. 2019, 9, 65. 10.3390/life9030065.

Szathmáry, E. Toward major evolutionary transitions theory 2.0. Proc Natl Acad Sci U S A. 2015, 112, 10104–10111.

Wiener, N. Cybernetics: Or Control and Communication in the Animal and the Machine. Cambridge (MA): MIT Press; 1948.

Wodak, S.J.; Paci, E.; Dokholyan, N.V.; et al. Allostery in its many disguises: from theory to applications. Structure 2019, 27, 566–578. 10.1016/j.str.2019.01.003.

Zimmermann, J.; Werner, E.; Sodei, S.; Moran, J. Pinpointing conditions for a metabolic origin of life: underlying mechanisms and the role of coenzymes. Acc. Chem. Res. 2024, 57, 3032–3043. 10.1021/acs.accounts.4c00423.

